# WNT/β-catenin increases 3D chromatin mobility and reorganisation during posterior cell state transitions

**DOI:** 10.64898/2026.10.05.756771

**Authors:** Annabelle A.M. Wurmser, Lingjun Mao, Nina Markevych, Gökçe G. Agsu, Yuze Cao, Ziwei Zhang, Mantas Jonaitis, Natasha De Winter, Benjamin D. Simons, Madapura M. Pradeepa, David Holcman, Thorsten E. Boroviak, David Klenerman, Shamith Samarajiwa, Srinjan Basu

## Abstract

Chromatin mobility, the exploration of 3D conformations by the chromatin fibre, is implicated in transcription and DNA repair, and is thought to decrease progressively as cells differentiate. Here, combining live-cell 3D single-nucleosome tracking with genomic approaches, we show that chromatin mobility instead changes in a cell-type-specific manner, increasing as pluripotent cells acquire posterior fates. This is accompanied by 3D genome reorganisation, including loss of Polycomb-associated interactions and emergence of CDX2-associated interactions. Withdrawal of posteriorising WNT/β-catenin signalling rapidly reduces chromatin mobility and partially restores Polycomb-associated chromatin interactions, while *Cdx1/2/4*-null cells fail to acquire these WNT/β-catenin-dependent changes. Together, our findings establish chromatin mobility as a regulated property of cell state, with WNT/β-catenin signalling and CDX factors acting as mediators that fine-tune the sampling of 3D chromatin interactions during cell state transitions.

## Main

The regulation of gene expression during development requires precise remodelling of 3D chromatin organisation, including of A/B compartments, topologically associating domains (TADs) and chromatin loops^1–7^. Yet how new chromatin conformations are explored during cell state transitions remains poorly understood. While chromosome conformation capture approaches have been instrumental in uncovering the widespread reconfiguration of chromatin interactions at increasingly high spatial resolution and complexity^8,9^, including at the single-cell level^10–13^, they fail to directly capture their dynamic nature^14^. Understanding chromatin remodelling during cell state transitions therefore requires considering both its 3D organisation and the biophysical properties that allow it to explore different conformations through space and time.

Polymer modelling and live-cell microscopy have established that chromatin behaves as a viscoelastic polymer whose physical properties depend on the spatial and temporal scales of observation^15–20^. While relatively stable over longer timescales (hours to days), on short temporal scales (seconds to minutes) TADs and chromatin loops are highly dynamic, short-lived, and display partial folding configurations, which can be stabilised by CTCF-cohesin complexes^21–23^. Consequently, chromatin is better described as a dynamic ensemble of conformations rather than as a single fixed state. This view also challenges simple associations between transcriptional activity, chromatin accessibility, compaction and mobility: active euchromatin can form condensed domains, while changes in chromatin mobility can occur independently of changes in compaction^24^. Functionally, chromatin conformational sampling modulates the duration and frequency, and thus probability, of interactions between genomic regions and nuclear proteins, including within chromatin domains previously considered relatively immobile, such as Polycomb domains and pericentromeric heterochromatin^25,26^. Consistent with this emerging view, changes in chromatin mobility have been implicated in various nuclear processes, including transcription, DNA replication, and DNA damage repair^27–37^.

Despite its emerging importance, how chromatin mobility is modulated during developmental cell state transitions remains largely unexplored. During these transitions, cells must rapidly integrate extracellular signals to make lineage decisions robustly while maintaining plasticity. These transitions are accompanied by extensive remodelling of genome organisation, yet how signalling pathways are translated into the physical dynamics of chromatin remains poorly understood. Modulating chromatin mobility could provide a rapid mechanism for reshaping the conformational landscape explored by the 3D genome, and hence transcriptional responses. Although some studies have reported greater chromatin mobility in pluripotent than in differentiated cells^16,34,38^, it remains unclear whether chromatin mobility changes progressively during differentiation or instead undergoes context-dependent regulation across distinct lineage transitions. Crucially, the molecular pathways that regulate chromatin mobility during development, and how these dynamic changes relate to higher-order genome organisation and transcriptional regulation, remain unknown.

Cell state transitions that occur during gastrulation and axis elongation provide a powerful model in which to address these questions. During these transitions, pluripotent epiblast cells integrate signalling cues, including WNT and FGF, to acquire a posterior identity and generate neuromesodermal competent progenitor cells able to differentiate into both spinal cord progenitors and somitic mesoderm^39,40^. These cell state transitions can be recapitulated *in vitro* and are accompanied by differential transcription factor binding alongside extensive remodelling of DNA accessibility and transcriptional programmes^41,42^. Importantly, the signalling pathways and downstream transcription factors controlling these transitions, including WNT/β-catenin and SOX2, T/Bra and CDX transcription factors, are well characterised, providing a tractable system to determine the interplay between developmental signals, chromatin mobility and organisation during cell state transitions.

Here, we combine live-cell 3D single-nucleosome tracking with genomic profiling to determine how chromatin mobility and organisation are coordinated during posterior cell state transitions. We find that chromatin mobility is dynamically regulated across developmental cell states through WNT/β-catenin signalling and the CDX1/2/4 transcription factor family. Together, our findings identify chromatin mobility as an additional layer of genome regulation that is modulated by developmental signals during lineage specification.

## Results

### Chromatin mobility increases as pluripotent cells transition toward neuromesodermal-competent progenitors and posterior spinal cord progenitors

To establish live-cell 3D single-molecule tracking (SMT) of single nucleosomes, we generated mouse embryonic stem cells (ESCs) stably expressing the core histone protein H2B fused with a C-terminal HaloTag (H2B-HaloTag) (**Fig. 1a**) and differentiated them using a well-characterised *in vitro* model of posterior axis elongation^39,41,42^. Naïve ESCs transition through primed epiblast-like cells (EpiLCs) and caudal epiblast-like cells (CEpiLCs), enriched for bipotent neuromesodermal-competent progenitors, before differentiating into spinal cord progenitors (SCPs) or somitic mesoderm (SM) (**Fig. 1b**). HaloTag labelling with the fluorophore JFX_554_ validated H2B-HaloTag expression (**Extended Data Fig. 1a**). Differentiation was validated by immunocytochemistry, with CEpiLCs co-expressing CDX2, SOX2 and T/BRA, and SCP and SM cells co-expressing SOX2/SOX1 and T/Bra/TBX6, respectively (**Extended Data Fig. 1b–c**).

**Figure 1.**
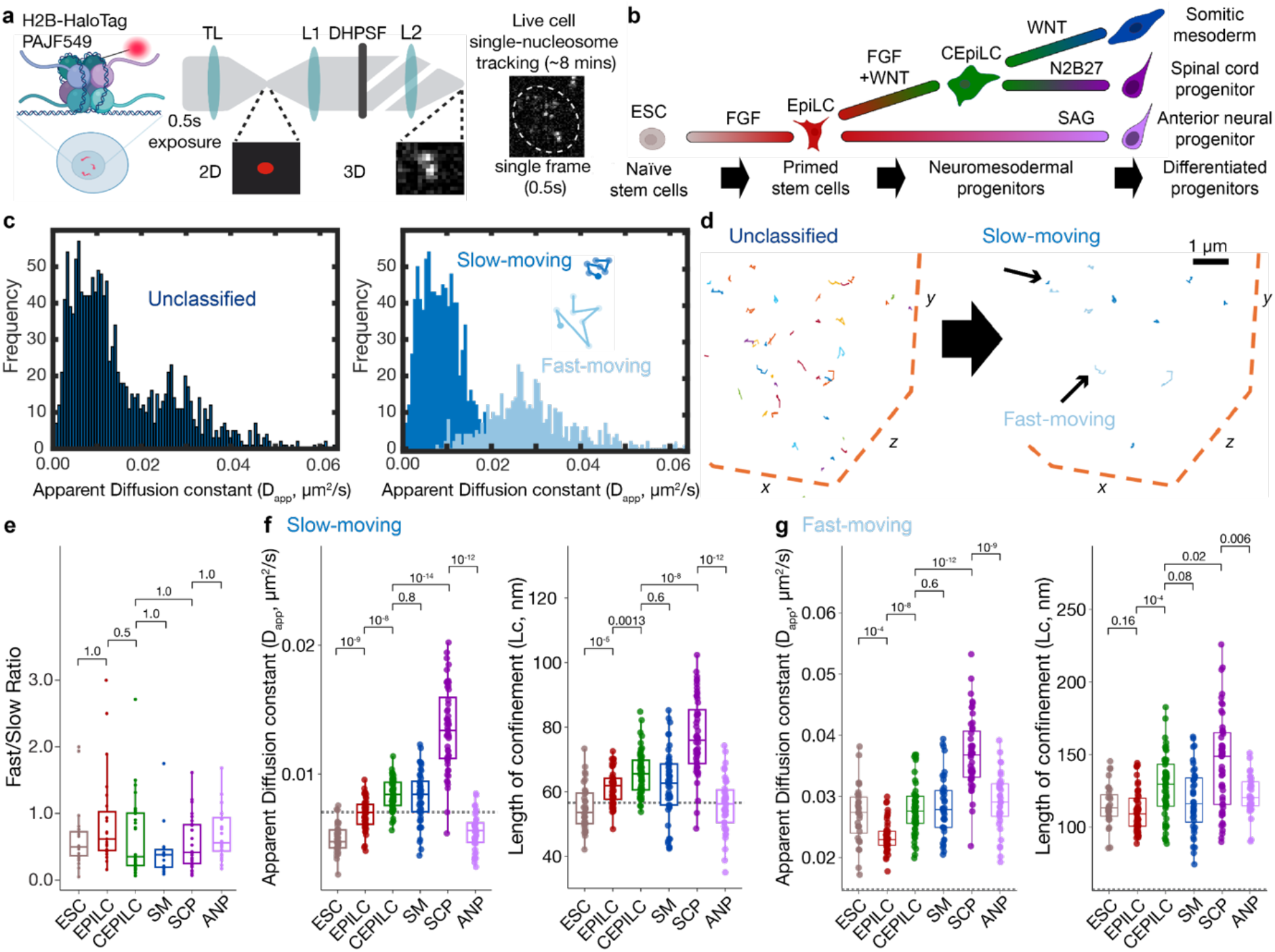
Chromatin mobility differs across cell types along the anterior–posterior axis. **(a)** Schematic of live single-nucleosome SMLM (single-molecule localisation microscopy) by use of a H2B-HaloTag fusion protein as part of a nucleosome structure with PA-JF_549_ Halo-ligand bound within the nucleus (left). Emission of the PA-JF_549_ fluorophore as a single point spread function is split in two lobes by the double helix point spread function (DHPSF) phase mask with the angle of rotation between the two lobes encoding the z-axis position (middle). Single frame from an SMLM movie with dotted lines showing overlaid nuclei including an inset zooming in for a detailed view of a single molecule, showing resolved double dots expected for a single molecule arising from a DHPSF phase mask (right). Created with BioRender.com. **(b)** Schematic of differentiation protocol: naïve pluripotent mouse embryonic stem cells (ESC) transition in FGF2 to primed pluripotent epiblast-like cells (EpiLCs) then in FGF2 and Chiron, which activates WNT/β-catenin signalling, to caudal epiblast-like cells (CEpiLCs) that contain bipotent neuromesodermal-competent progenitors. CEpiLCs then differentiate into either spinal cord progenitors (SCPs) in N2B27 alone, somitic mesoderm (SM) in CHIRON or anterior neural progenitors (ANPs) in SAG. **(c)** Histograms showing average apparent diffusion constant (D_app_) extracted per sliding window (9 frames, 4.5 s) of a trajectory (left) before classification and (right) after classification into fast-moving (light blue) or slow-moving chromatin (dark blue) using a Gaussian mixture model. **(d)** Example trajectories (left) pre-classification and (right) post-classification for an EpiLC sample with nucleus border depicted as orange dotted line. Pre-classification trajectories are shown as different colours per trajectory. Post-classification trajectories are defined as fast-moving (light blue) or slow-moving chromatin (dark blue). **(e)** Boxplot showing no significant difference in the ratios of fast-to slow-moving chromatin across cell types. Statistical significance was determined by Kruskal–Wallis test, followed by Dunn’s post-hoc test with Holm correction for multiple comparisons (*n* > 38 FOVs). **(f-g)** Boxplots show the apparent diffusion constant (D_app_), and confinement radius (Lc) of **(f)** slow-moving and **(g)** fast-moving chromatin trajectories, averaged per field of view (FOV). Colours indicate cell types labelled in (a). Data from 5-10 FOVs (≥5 cells per FOV) per technical replicate (≥2) for each biological replicate (≥3). Statistical significance was calculated using Dunn’s post-hoc test with Benjamini–Hochberg correction for multiple testing (*n* > 38 FOVs). The black dotted line depicts the value from dye on a coverslip to indicate where measurements are below the detection limit of the microscope.

**Extended Data Figure 1.**
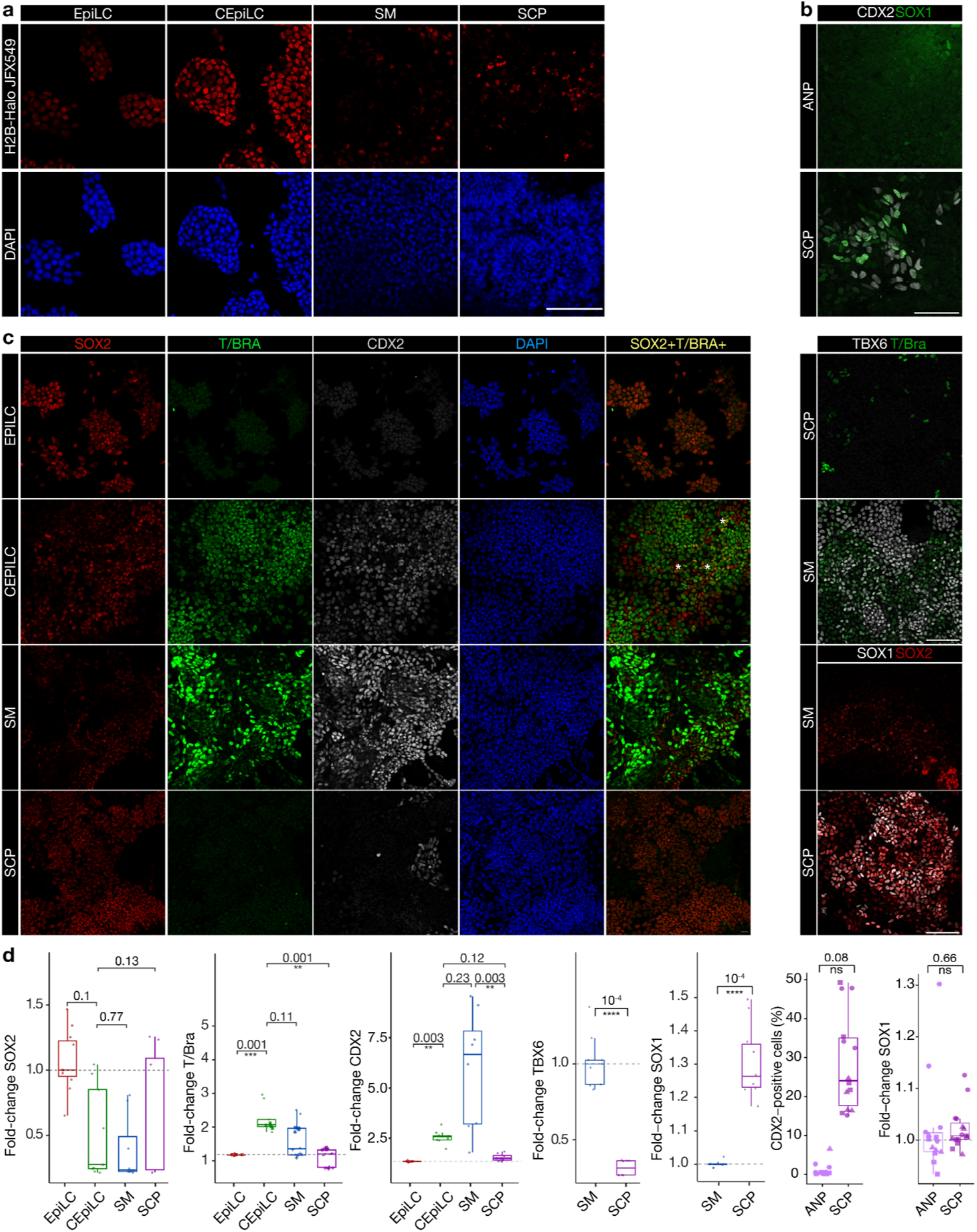
Validation of H2B-Halo expressing mESCs. **(a)** Representative fluorescence microscopy images showing nuclear JFX_554_ signal (top row, red) and DAPI staining (bottom row, DAPI) confirm H2B-HaloTag fusion protein expression throughout differentiation. Scale bar 100 μm. **(b)** Representative fluorescence microscopy images showing nuclear TF signal in indicated cell types. Scale bar 100 μm. **(c)** Box plot showing fold-change in TF expression per sample relative to reference. For each individual cell, fluorescence intensity was normalised to DAPI. The mean expression was calculated for each FOV (≥50 cells/FOV). Statistical significance was assessed using a Kruskal–Wallis test followed by Dunn’s multiple-comparisons test, with p-values adjusted using the Benjamini– Hochberg method. Percentage of CDX2-positive cells was based on the 99th percentile highest expression in ANPs. Statistical comparison between biological replicates was carried out using a Wilcoxon-rank test. A binomial GLMM using positive/negative cell counts per FOV with biological replicate as a random intercept estimated higher odds of CDX2 positivity in SCPs (OR = 40.7, 95% CI 20.5–80.5, P = 2.2 × 10^−26^). ([SOX2/CDX2/T – EpiLCs: n=2324 cells; CEpiLCs: 4115 cells; SM: n=4541; SCP: n= 3642; TBX6 – SM: n=6813, SCP: n=7205; SOX1 – SM: n= 7776; SCP: n=7425; CDX2/SOX1 – ANPs: n =7923 cells; SCPs: n=3991 cells] across >2 biological replicates and >8 technical replicates).

**Extended Data Figure 2.**
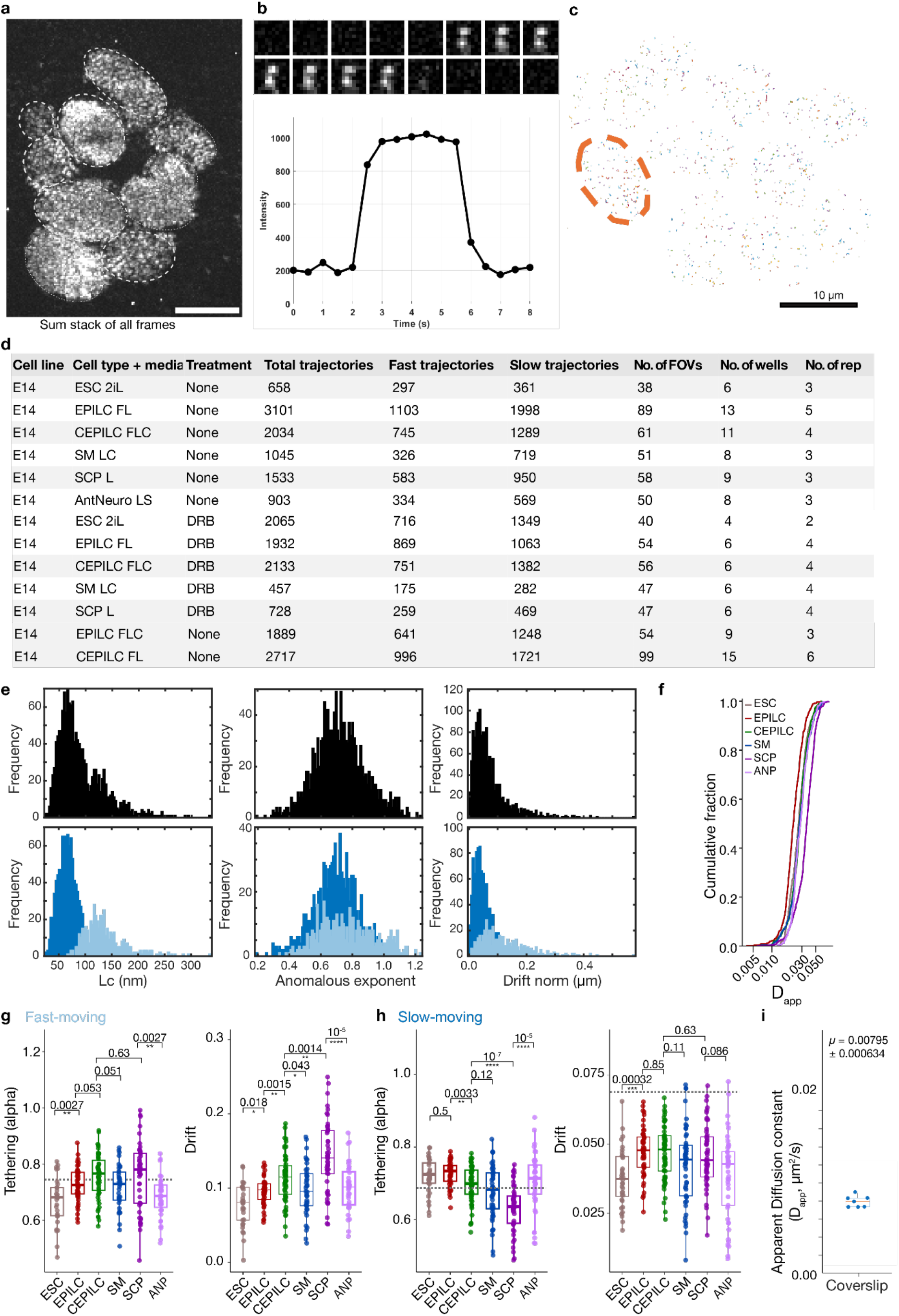
Quality control for SMLM. **(a)** Sum stack of all frames showing individual double dots within nuclei. Scale bar: 10um. **(b)** (Top) Frame montage showing a single H2B molecule over the course of 7.5 s (16 frames), with signal lasting 3.5s (7 frames). (Bottom) Intensity profile over time for molecule above. **(c)** Example trajectories from a field of view (FOV) showing approximately 10 nuclei with orange dotted line delineating nucleus shown in Fig. 1a. **(d)** Table showing number of unclassified and classified trajectories, FOVs, wells and biological replicates per dataset. **(e)** Histograms showing confinement radius (Lc), drift (D_norm_) and anomalous exponent (tethering, α) extracted per sliding window (9 frames, 4.5 s) of a trajectory (top) before classification and (bottom) after classification into fast-moving (light blue) or slow-moving chromatin (dark blue) using a Gaussian mixture model. **(f)** Empirical Cumulative Distribution Function (ECDF) of single-trajectory D_app_ values in WT cells across the indicated developmental states. All trajectories from FOVs retained after FOV-level 1.5×IQR outlier filtering are shown. ECDFs are descriptive; statistical inference was performed using FOV-level summary values in (Fig. 1e). **(g-h)** Boxplots show the tethering (anomalous exponent ɑ) and drift (D_norm_) of **(g)** fast-moving and **(h)** slow-moving/immobile chromatin trajectories, averaged per FOV. Data from 5-10 FOVs (≥5 cells per FOV) per technical replicate (≥2) for each biological replicate (≥2). Statistical significance assessed using Wilcoxon-rank tests with Benjamini–Hochberg correction for multiple testing. The grey box depicts the precision limit where measurements are below the detection limit of the microscope. **(i)** Boxplot showing D_app_ of dye on a coverslip (n = 7).

For live-cell 3D SMT, H2B-HaloTag was labelled with the bright photoactivatable fluorophore PA-JF_549_ (**Fig. 1a**)^43^. Cells were imaged using a double-helix point spread function microscope enabling 3D tracking across a 4-μm axial range^44–46^. Imaging at 500-ms resolution substantially reduces the detection of freely diffusing molecules through motion blurring, enabling measurement of global chromatin mobility^44,46–48^. Single H2B molecules were detected at high signal-to-noise and exhibited single-step photobleaching, a hallmark of single-molecule detection (**Extended Data Fig. 2a,b**).

To monitor global chromatin mobility, individual single-molecule localisations were assembled into trajectories (**Fig. 1c-d**) and classified into both slow- and fast-moving sub-diffusive chromatin states based on a sliding window of 9 consecutive images (4.5 seconds) using our previously described four-parameter SMT classification pipeline (**Fig. 1c,d, Extended Data Fig. 2c-e**)^30,44,46,49^. At this time and length scale, chromatin mobility does not change during the cell cycle^50^. The proportion of fast- and slow-moving chromatin populations did not significantly change across cell states (**Fig. 1e, Extended Data Fig. 2f**). We therefore compared four biophysical parameters from each motion class: apparent diffusion coefficient (D_app_), anomalous exponent (α; reflecting chromatin tethering), confinement length (Lc; the radius explored by chromatin) and normalised drift magnitude (D_norm_; reflecting directionality) (**Fig. 1f,g, Extended Data Fig. 2g,h**). In ESCs, slow- and fast-moving chromatin exhibited D_app_ values of 0.005±0.001 μm^2^s^−1^ and 0.027±0.005 μm^2^s^−1^ respectively. The D_app_ of the fast-moving chromatin state decreased as ESCs transitioned to EpiLCs, then continued to increase during posteriorisation and differentiation into SCPs, but not as CEpiLCs differentiated into SM (**Fig. 1f,g**). Consistent trends were observed for Lc, α and D_norm_ (**Fig. 1f,g, Extended Data Fig. 2g,h**), where Lc decreased slightly from ESCs to EpiLCs, increased significantly from EpiLCs to CEpiLCs, and was highest in SCPs, while remaining comparable in CEpiLCs and SM. Meanwhile, the D_app_ of the slow-moving chromatin state was below the detection limit of the microscope, indistinguishable from that of immobile dye molecules on a coverslip, which had a residual D_app_ of 0.008±0.001 μm^2^s^−1^ (63 ± 3 nm localisation precision) (**Extended Data Fig. 2i**).

These findings contrast with a previous study which reported lower chromatin mobility in anterior neural progenitors (ANPs) than in ESCs^38^. We therefore measured chromatin mobility during anterior neuroectoderm differentiation from EpiLCs in the presence of SAG, a Sonic Hedgehog agonist^41^. We found no significant differences relative to ESCs but a significant decrease relative to SCPs (**Fig. 1f,g, Extended Data Fig. 2g,h**). Thus, our findings suggest that chromatin mobility varies across both cell states and the anterior–posterior axis, with higher mobility in CEpiLCs than EpiLCs and in posterior SCPs than ANPs.

### A/B compartment segregation and TAD size increase during posteriorisation

To relate posterior-specific chromatin mobility changes to 3D chromatin organisation, we used Micro-C^51^ to map genome-wide 3D chromatin organisation at 10 kb resolution (**Fig. 2a, Extended Data Fig. 3a**). The genome was partitioned into megabase-scale A and B compartments, associated with gene-rich/euchromatic and gene-poor/heterochromatic regions, respectively^52^. The data revealed an increase in compartment strength during posteriorisation, attributed primarily to increased B-B interactions, with little change observed during subsequent CEpiLC-to-SM or CEpiLC-to-SCP differentiation (**Extended Data Fig. 3b**). Compartment switching was also greatest during posteriorisation: 9.5% of genomic bins switched from A to B and 4.9% from B to A, while 85.6% retained their compartment identity (**Fig. 2b,c, Extended Data Fig. 3c**). In contrast, 3.2% and 3.9% of bins switched from A to B and 4.2% and 4.0% from B to A during CEpiLC-to-SM and CEpiLC-to-SCP differentiation, respectively. Thus, posteriorisation is accompanied by increased segregation of A/B compartments and extensive reassignment of compartment identity, consistent with large-scale reorganisation observed during anterior neural lineage specification^53^.

**Figure 2.**
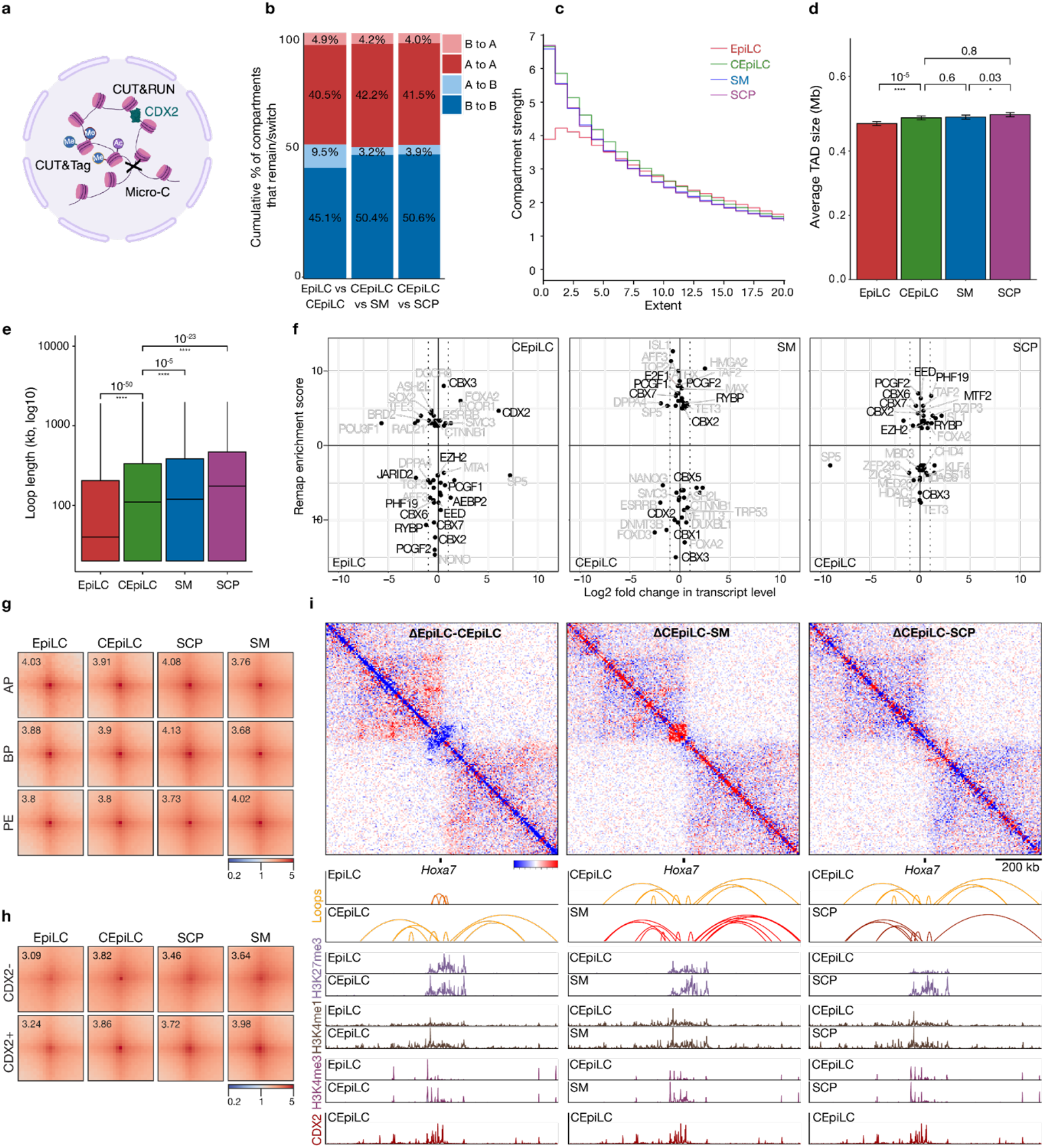
Major 3D genome reorganisation occurs during posteriorisation correlating with a loss of loops associated with Polycomb-associated bivalent promoters and a gain of loops linked to CDX2. **(a)** Schematic of experimental approach. Micro-C captures 3D chromatin interactions via MNase digestion and proximity ligation. CUT&Tag profiles histone modifications using antibody-targeted Tn5 tagmentation. CUT&RUN maps CDX2 occupancy using antibody-targeted pAG–MNase digestion. Made with Biorender. **(b)** Quantification of A/B compartment switches between cell types, expressed in percentages at 1Mb resolution for each cell state transition. **(c)** Line plot showing compartment strength at different values of the extent of interaction across cell types, calculated as the ratio (AA + BB)/(AB + BA) from saddle plots of interactions between A and B compartments (see **Extended Data Figure 3b**), where higher values indicate stronger compartmentalisation. **(d)** Bar plot of the average TAD size in Mb (million base pairs), with error bars representing 95% confidence interval. Statistical testing using a Mann-Whitney-Wilcoxon two-sided test with Benjamini-Hochberg correction. **(e)** Box and whisker plot of loop length per cell type for high-confidence loops identified by FitHiC2 (FDR<10^−5^). Statistical testing using a Mann-Whitney-Wilcoxon two-sided test with Benjamini-Hochberg correction. **(f)** Correlation plot of log2-fold change in transcript levels (bulk RNA-seq) and the ReMap enrichment score (Micro-C) reveals a functional role for CDX2 or Polycomb (highlighted in black). The dotted line corresponds to log2-fold change > 1. Only significant DEGs are plotted. **(g)** APA plots of CRE classes by cell type. (**h**) APA plots of loops with and without CDX2 bound at one or both of the loop anchors. (**i**) Micro-C differential contact maps and epigenomic profiles at the *Hoxa7* locus (chromosome 6, 52.19 Mb). Top, differential contact maps comparing EpiLC and CEpiLC (left), CEpiLC and SM (middle), and CEpiLC and SCP (right). Red and blue indicate relative gains and losses in chromatin domains, respectively. Bottom, corresponding chromatin loops, H3K27me3, H3K4me1, and H3K4me3 CUT&Tag tracks and CDX2 CUT&RUN tracks for the indicated cell types. Scale bar, 200 kb.

**Extended Data Figure 3.**
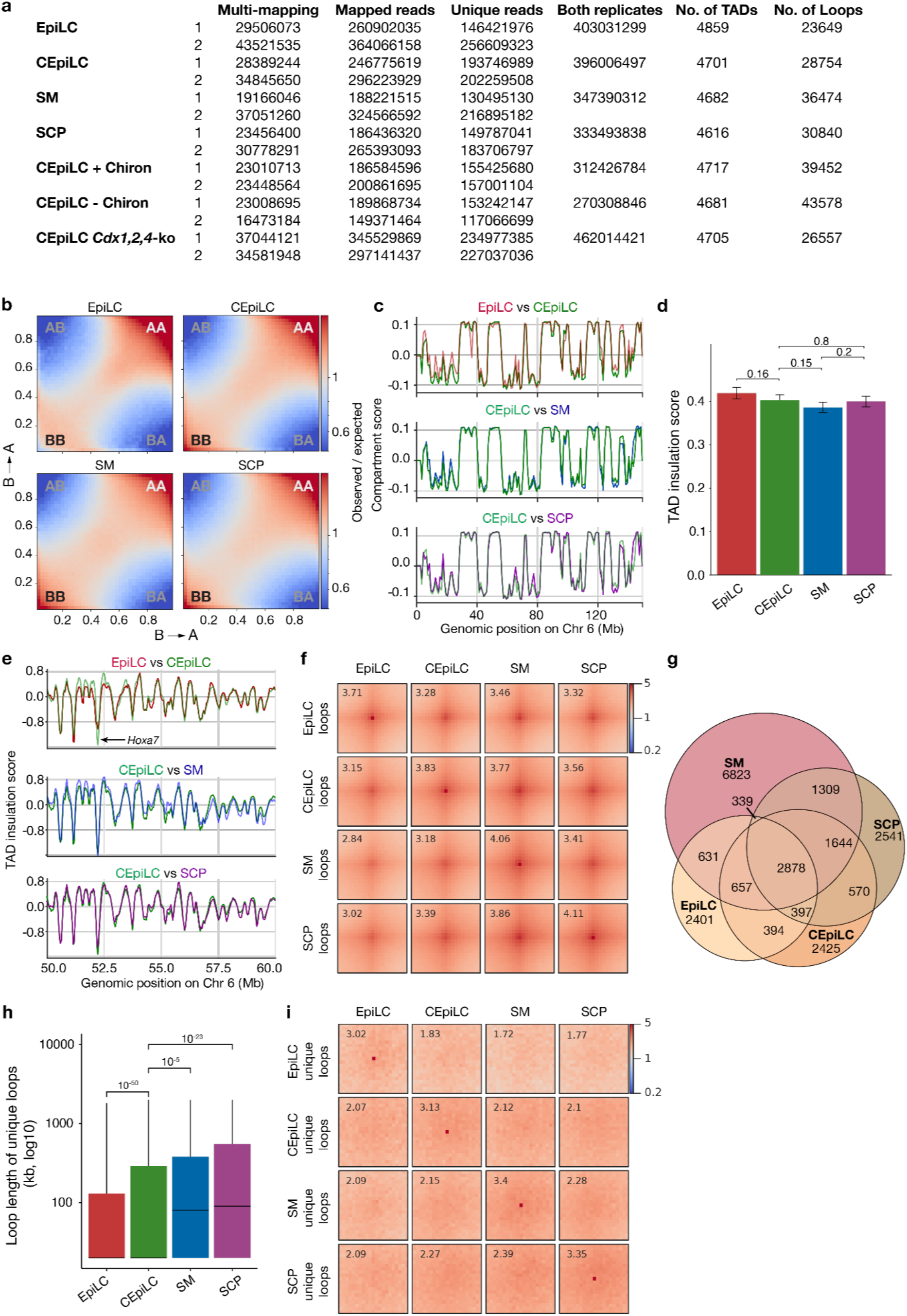
Micro-C data reveals chromatin reorganisation during posteriorisation. **(a)** Summary table of Micro-C sequencing metrics, including total, mappable and uniquely mapped reads, valid 3D contacts, numbers of A/B compartments, topologically associating domains (TADs) and significant chromatin loops identified by FitHiC2 (q < 10⁻⁵). **(b)** Genome-wide saddle plots derived from Micro-C contact matrices for EpiLC, CEpiLC, SM and SCP cells showing compartment strength across differentiation. Genomic bins were ranked according to compartment eigenvector (PC1) values and divided into equal quantiles from B to A compartment identity (x-axis) and A to B compartment identity (y-axis). Colours represent observed/expected contact enrichment between compartment quantiles, with red indicating enriched interactions and blue indicating depleted interactions. Increased enrichment of A-A and B-B interactions, together with depletion of A-B interactions, indicates stronger compartmentalisation. **(c)** Line plot of the compartment score profiles across chromosome 6 (chr6) for EpiLC, CEpiLC, SM and SCP cells, showing A compartments as positive values and B compartments as negative values. **(d)** Bar plot of the absolute TAD insulation score for EpiLC, CEpiLC, SM and SCP cells. Error bars represent 95% confidence intervals. Statistical significance was assessed using two-sided Wilcoxon tests with Benjamini–Hochberg correction. **(e)** Line plot of the TAD insulation score calculated per cell type with pairwise comparisons highlighting genomic regions undergoing TAD changes at the chromosome 6 *Hox* region (*Hoxa7* locus at 52.19 Mb). **(f)** Aggregate peak analysis (APA) plots showing the enrichment of all chromatin loops identified in EpiLC, CEpiLC, SM and SCP cells across all cell types. **(g)** Euler diagram showing the overlap between significant chromatin loops detected during multiway comparisons of all cell types. **(h)** Box and whisker plot showing distribution of genomic distances spanned by chromatin loops unique to EpiLC, CEpiLC, SM and SCP cells. **(i)** APA plots of cell-type-specific loops across all cell types, demonstrating that loops identified as cell-type-specific exhibit their strongest aggregate enrichment in the corresponding developmental state.

**Extended Data Figure 4.**
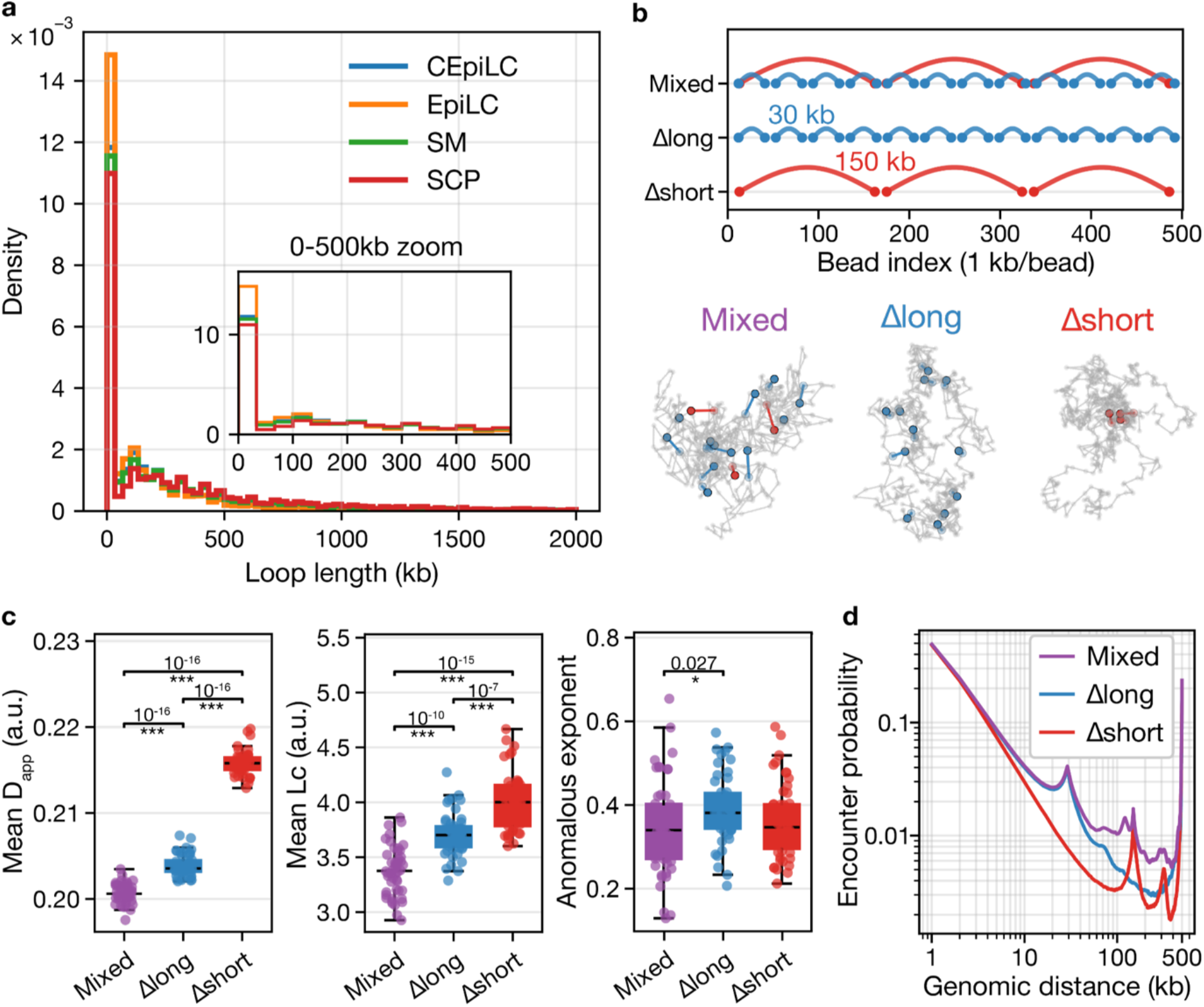
Polymer simulations uncover how loop size distribution influences chromatin mobility and the encounter probability of chromatin interactions at various genomic distances. **(a)** Density distribution of loop length in kb across cell types. Inset shows an enlarged view of the distribution between 0 and 500 kb. **(b) Top:** Schematic representation of the polymer model, illustrating three simulation conditions including a mixed population of short and long loops (Mixed), selective loss of long loops (Δlong) and selective loss of short loops (Δshort)**. Bottom:** Representative polymer realisations for each condition. Grey tracts represent the polymer backbone, coloured beads represent loop anchors, and coloured lines represent loop-forming harmonic bonds (red, 150 kb long loops; blue, 30 kb short loops). **(c)** Comparison of per-polymer mean apparent diffusion constant (D_app_), length of confinement (Lc), and anomalous exponent (α) across the three simulation conditions. [Mann-Whitney-Wilcoxon two-sided test with Benjamini-Hochberg correction for multiple testing, *n* = 50 realisations per condition] **(d)** Encounter probability of two loci as a function of genomic distance (kb) across the simulated conditions.

**Extended Data Figure 5.**
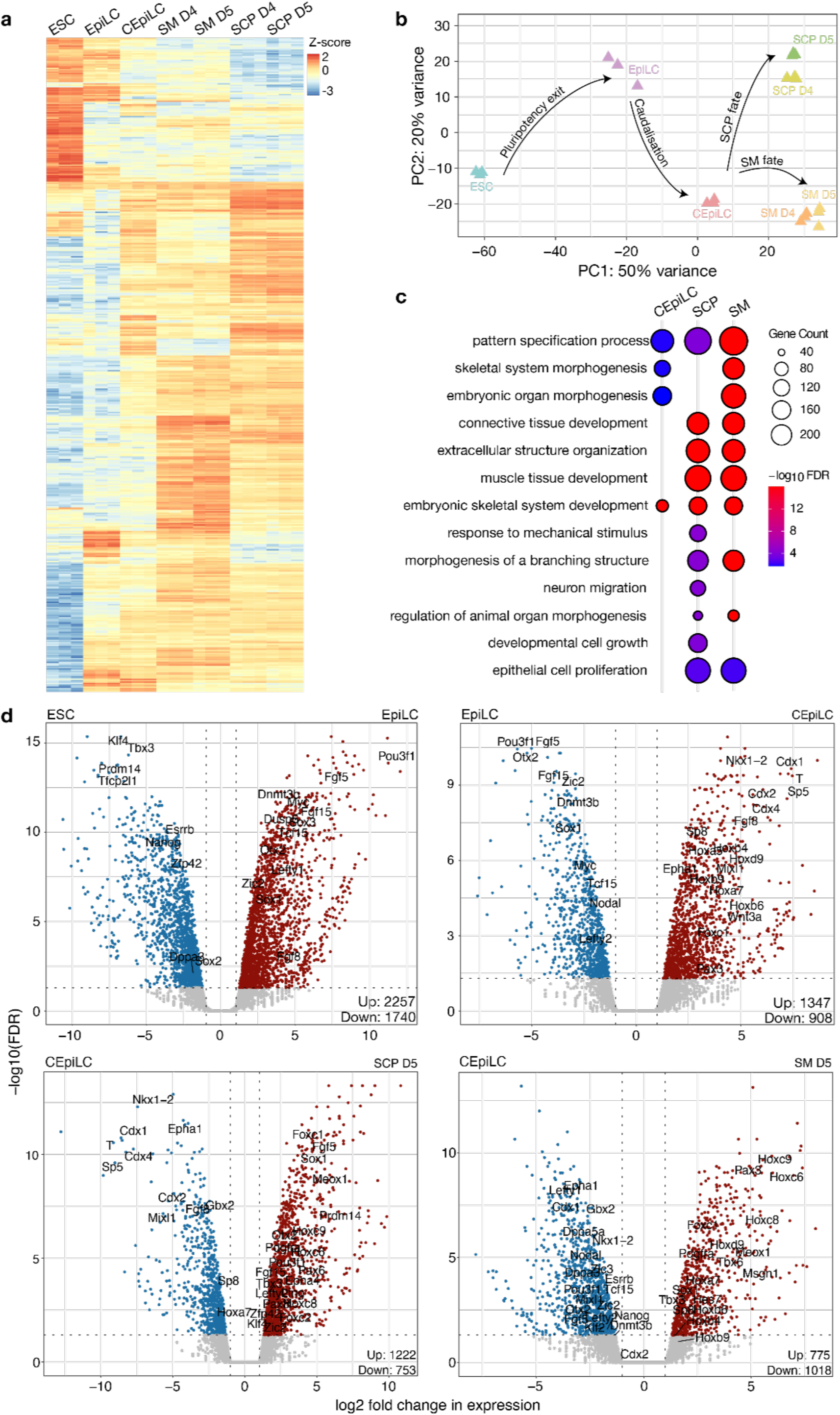
Quality control of RNA-seq datasets. **(a)** Heatmap showing relative expression of cell type-specific marker genes. Scale indicates z-scores. **(b)** Principal component analysis of RNA-bulk seq transcriptomes across differentiation stages shows cell type-specific clustering. Points represent individual technical replicates, with each coordinate based on global gene expression profiles (counts normalised and variance-stabilised). Arrows illustrate inferred transcriptional trajectories: pluripotency exit, posteriorisation, SCP fate acquisition, and SM fate acquisition. Percentages on the axes indicate the proportion of variance explained by each principal component. **(c)** Dot plot of relevant molecular gene ontology (GO) terms enriched at each time point relative to EpiLCs. Dot colour scale representing log_10_(FDR) value. Dot diameter representative of the number of genes at a given time point that overlap with the respective GO term. **(d)** Volcano plots of differentially expressed genes (DEGs) across time points.

**Extended Data Figure 6.**
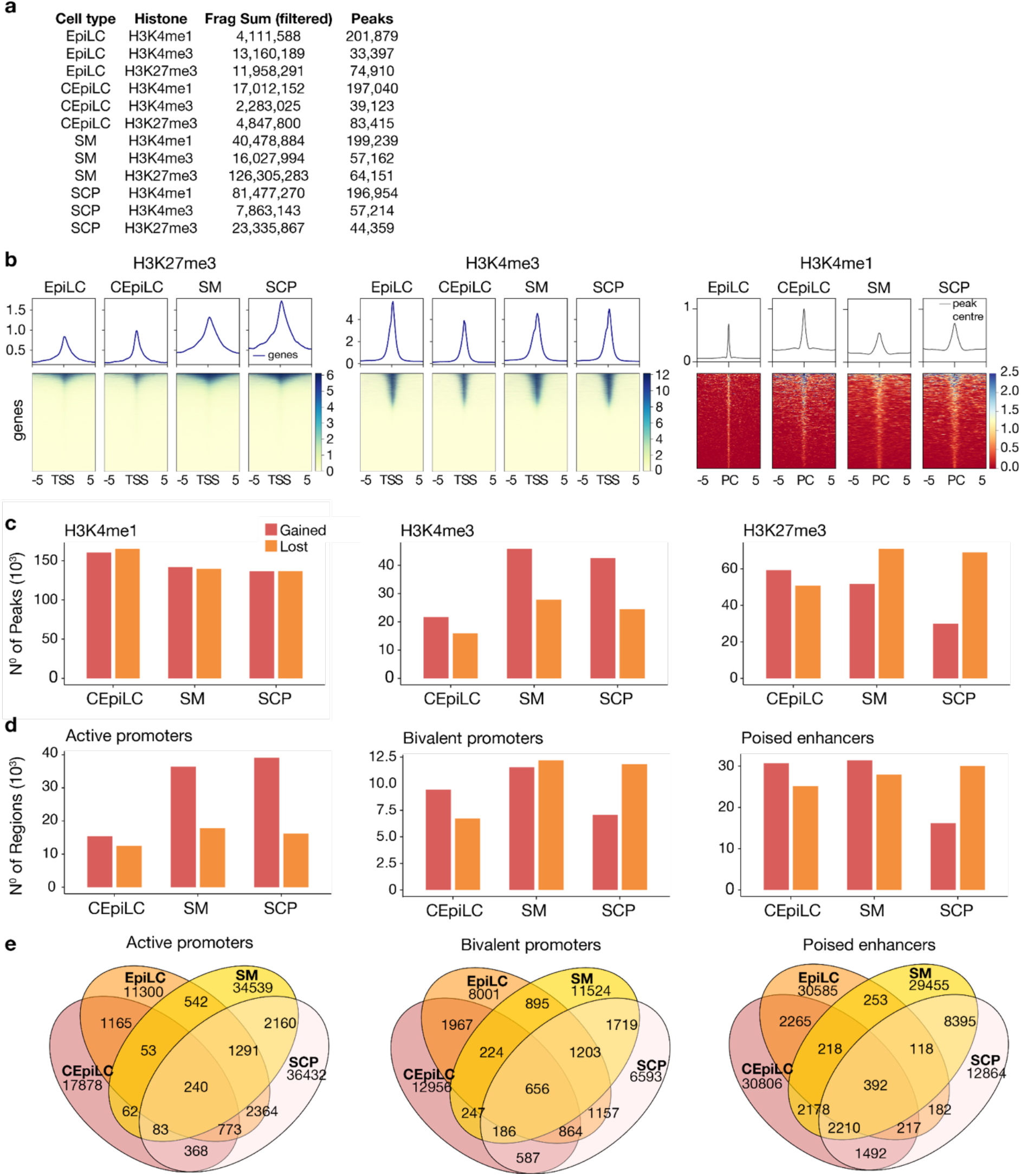
CUT&Tag datasets reveal global redistribution of histone marks at cis-regulatory elements during posteriorisation and differentiation. **(a)** Number of unique reads and identified peaks from CUT&Tag of corresponding cell types and histone marks. **(b)** Averaged signal profiles (above) and heatmaps (below) for H3K4me3 and H3K27me3 at 5 kb either side of annotated transcription start sites (TSS). **(c)** Number of histone peaks lost and gained relative to the preceding cell type. **(d)** Number of regions per cis-regulatory element class lost and gained relative to the preceding cell type. **(e)** Venn diagrams showing unique and shared loops from multiway comparison of loops during cell state transitions for bivalent and active promoters (left) and poised enhancers (right).

**Extended Data Figure 7.**
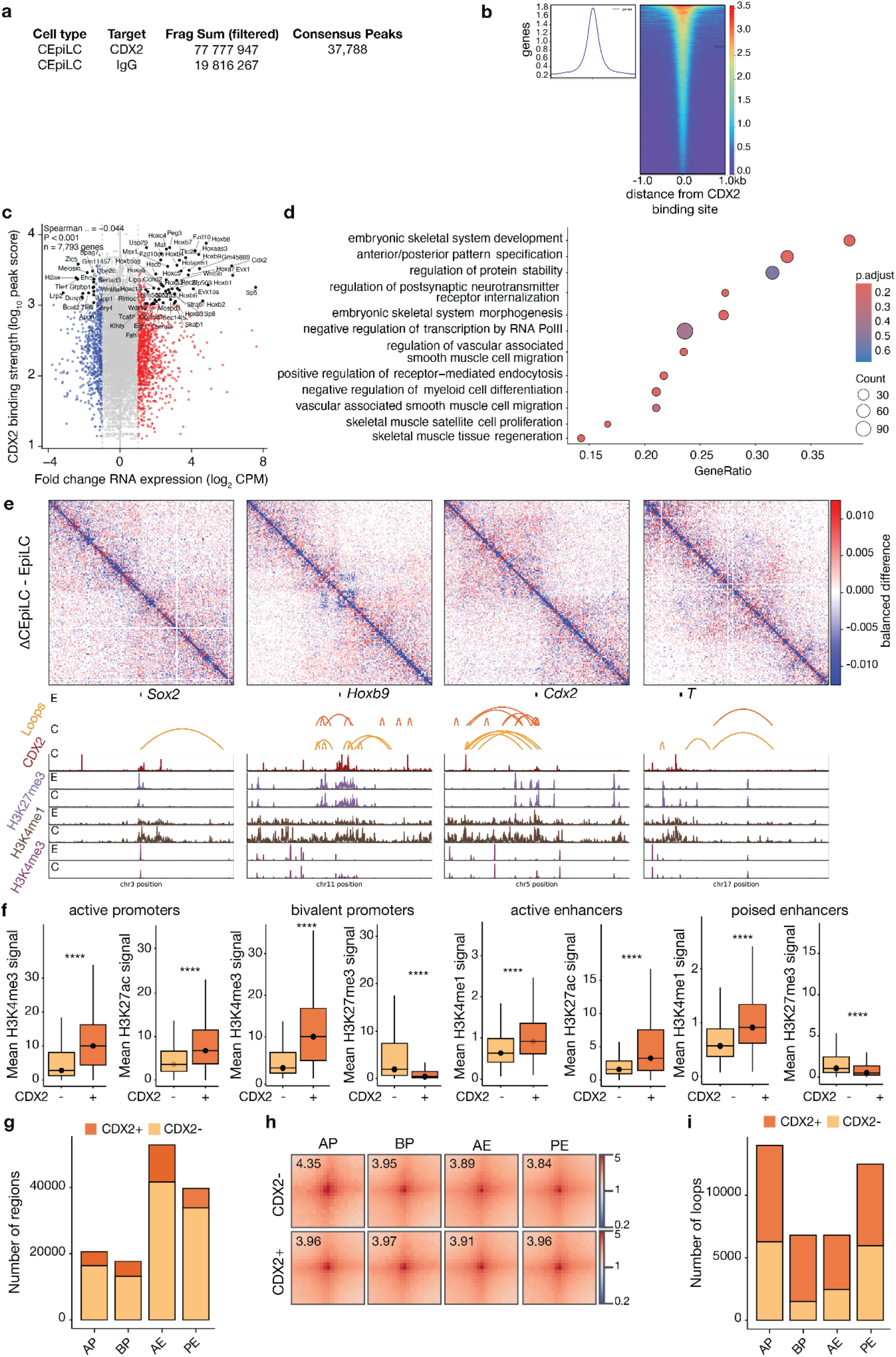
CUT&RUN and CUT&Tag datasets reveal changes in CDX2 binding and global redistribution of H3K4me3 and H3K27me3 histone marks at cis-regulatory elements during posteriorisation and differentiation. **(a)** Table of filtered reads and MACS consensus peaks per condition. **(b)** Heatmap showing CDX2 signal 5 kb either side of TSSs across the genome. **(c)** Scatter plot of CDX2 binding strength versus EpiLC-to-CEpiLC RNA expression change. Genes in the top 10% for both binding strength and absolute expression change are labelled. Spearman’s ρ=−0.044, P<0.001, n=7,793. **(d)** GSEA analysis of CDX2-bound promoters. **(e)** Representative loci showing chromatin loops, histone marks and CDX2 binding during the EpiLC-to-CEpiLC transition. **(f)** Mean histone signal at CDX2 bound and control regions for the different CRE classes in CEpiLCs. **(g)** Bar plot of the number of regions by CDX2 binding status for the different CRE classes. **(h)** APA plots showing loop strength according to CDX2 binding for all CRE classes **(i)** Bar plot of the number of loops according to anchor overlap with CDX2 binding.

We next examined sub-megabase-scale TADs formed by CTCF-cohesin loop extrusion^54–59^. Global TAD insulation scores did not significantly change (**Extended Data Fig. 3d**). However, TAD size significantly increased during posteriorisation (**Fig. 2d**). At the *Hox* locus on chromosome 6, posteriorisation coincided with a local decrease in insulation score (**Fig. 2i, Extended Data Fig. 3e**), indicative of strengthened TAD insulation at a locus known to undergo dynamic reorganisation during anterior–posterior patterning^60^.

Together, these results indicate that posteriorisation is accompanied by strengthening and reassignment of A/B compartments alongside increased TAD size, coinciding with increased chromatin mobility (**Fig. 1**). However, chromatin mobility increased further during SCP specification despite more limited compartment switching, arguing against mobility arising solely from compartment switching. Moreover, increased mobility during posteriorisation coincided with expansion and increased connectivity of the B compartment, indicating that enhanced chromatin mobility was not restricted to transcriptionally active A compartments.

### Switch from Polycomb-associated to CDX2-associated chromatin loops during posteriorisation

Having observed a poor relationship between chromatin mobility and compartment-level changes, we next examined chromatin organisation at finer genomic scales. We generated local contact matrices at 10 kb resolution and identified high-confidence chromatin loops using FitHiC2 (**Extended Data Fig. 3a,f**)^61^. This revealed extensive reorganisation of loops across differentiation (**Extended Data Fig. 3g-i**). Interestingly, loop length increased significantly during posteriorisation and further during differentiation towards SM and SCP identities, with the longest loops observed in SCPs (**Fig. 2e**). These loop lengths were broadly consistent with the progressive changes in chromatin mobility previously observed (**Fig. 1f-g**), prompting us to investigate whether alterations in loop architecture could account for the observed differences in mobility. Given the bimodal distribution of loop lengths and a shift towards longer loops in CEpiLCs and SCPs **(Extended Data Fig. 4a)**, we performed polymer simulations to determine whether the loss of short loops or the gain of long loops would best recapitulate the observed chromatin mobility changes **(Extended Data Fig. 4b)**. This revealed that, while a loss of short-range loops led to a significant increase in chromatin mobility **(Extended Data Fig. 4c)**, the gain of long-range loops had only a minimal impact on chromatin mobility but increased the likelihood of encounters between distal chromatin regions **(Extended Data Fig. 4d)**. Together, this suggested a possible link between chromatin mobility and the reorganisation of local chromatin interactions during differentiation.

To identify candidate regulators associated with this reorganisation, we intersected cell type-specific loop anchors with experimentally validated DNA-binding sites of nuclear proteins from the ReMap 2022 ChIP-seq atlas and ranked proteins by their relative enrichment across cell states^44,62^. EpiLC-specific loops were predominantly associated with Polycomb group (PcG) proteins, including the Polycomb Repressive Complex 1 (PRC1) subunits CBX2, CBX6, CBX7, RYBP and PCGF2, the Polycomb Repressive Complex 2 (PRC2) components PHF19, EZH2 and EED, and the PRC2 cofactor AEBP2 (**Fig. 2f**). This is consistent with previous reports demonstrating preferential association of Polycomb proteins with chromatin loops connecting bivalent promoters in primed pluripotent cells^44,63,64^. Posteriorisation was accompanied by a marked shift in the proteins associated with chromatin loops. Loop anchors became depleted for Polycomb-associated proteins and instead showed enrichment for transcription factors characteristic of the caudal epiblast state, including SOX2 and the WNT-responsive factors CDX2 and β-catenin (CTNNB1) (**Fig. 2f**). Heterochromatin Protein 1γ (CBX3) was also enriched, consistent with the increased B-B compartment interactions observed during posteriorisation (**Extended Data Fig. 3b**).

As cells acquired SM and SCP fates, Polycomb-associated proteins, including PCGF2, CBX2, CBX6/7, RYBP and EED, became enriched again, suggesting their reduction was transient in the CEpiLC state (**Fig. 2f**). Lineage-specific associations also emerged: SCP-specific loops were enriched for the neural-specifying transcription factor FOXA2, whereas SM-specific loops were enriched for the DNA demethylase TET3, previously implicated in mesoderm differentiation^65^. Thus, posteriorisation represents a transient shift from Polycomb-associated towards WNT-responsive chromatin interactions, followed by the emergence of lineage-specific regulatory interactions in the more differentiated cell states.

To determine whether changes in these loop-associated factors were accompanied by changes in their expression level, we performed bulk RNA sequencing across differentiation (**Extended Data Fig. 5a,b**). Differential expression and Gene Ontology analyses confirmed the expected progression from pluripotency through posteriorisation to lineage specification (**Extended Data Fig. 5c,d**). Notably, factors associated with two major developmental regulatory systems emerged from the comparison of loop-anchor enrichment and gene expression (**Fig. 2f**). The Polycomb-associated factors RYBP and JARID2 were differentially expressed and enriched at EpiLC-specific loop anchors, whereas CDX2 was strongly upregulated during the EpiLC-to-CEpiLC transition and among the highest-ranked proteins at CEpiLC-specific loop anchors. More broadly, changes in protein-associated chromatin interactions were not necessarily driven by corresponding changes in expression levels, suggesting additional regulation at the level of genomic occupancy.

Collectively, our findings suggest that exit from pluripotency, posteriorisation and subsequent lineage specification are accompanied by extensive remodelling of 3D chromatin organisation involving both chromatin regulators and lineage-specifying transcription factors. The increase in chromatin mobility during posteriorisation coincided with a temporal reduction in Polycomb-associated chromatin interactions alongside an increase in CDX2-associated interactions. However, Polycomb-associated interactions were enriched again in more differentiated SM and SCP states, without a corresponding loss in chromatin mobility. These observations indicate that regulatory mechanisms beyond that of Polycomb contribute to the increase in chromatin mobility during posterior fate specification.

### Posteriorisation is accompanied by transient remodelling of PcG-associated and CDX2-bound regulatory elements

Given the transient loss of Polycomb-associated proteins and emergence of CDX2 during posteriorisation, we examined how Polycomb-regulated and CDX2-bound cis-regulatory elements (CREs) are remodelled **(Fig. 2a)**. As Polycomb-associated proteins occupy CREs including bivalent promoters and poised enhancers^66,67^, we annotated active promoters (H3K4me3), bivalent promoters (H3K4me3 and H3K27me3) and poised enhancers (H3K4me1 and H3K27me3) using CUT&Tag (**Extended Data Fig. 6a,b**)^68^. Quantification of H3K27me3-, H3K4me1- and H3K4me3-associated peaks revealed dynamic gains and losses between cell types (**Extended Data Fig. 6c-e**), yet these changes did not correlate with the transient loss of Polycomb-associated loops (**Fig. 2g**). Thus, the transient reorganisation of Polycomb-associated loops is not simply explained by changes in H3K27me3 enrichment.

We therefore asked whether the newly established chromatin interactions were preferentially associated with CDX2 occupancy assayed by CUT&RUN (**Extended Data Fig. 7a,b**). Consistent with the established role of CDX2 in specifying posterior identity in CEpiLCs, CDX2 occupied promoters associated with WNT-responsive developmental genes, including *Wnt8a*, *Hoxa1* and *Hoxb9*, and bound its own promoter, consistent with recently described autoregulatory loops (**Fig 2i; Extended Data Fig. 7c,d**)^69^. CDX2-bound promoters were enriched for developmental processes including somitogenesis, WNT signalling, anterior– posterior patterning, muscle development and neurogenesis (**Extended Data Fig. 7e**). Together, these findings link CDX2 binding with posterior development.

Having established that CDX2 binds promoter elements of posterior developmental genes, we next examined how its occupancy relates to chromatin interactions during posteriorisation. Aggregate peak analysis demonstrated that CDX2-bound CEpiLC-specific loops progressively strengthened during posteriorisation and further increased during SM differentiation (**Fig. 2h)**. Because we also detected CDX2 binding at enhancers and bivalent promoters associated with key developmental regulators, including *T/Bra* and *Sox2* (**Extended Data Fig. 7c**), we next asked whether the strengthening of CDX2-associated loops was accompanied by changes in local histone modifications at CREs. CDX2-bound bivalent promoters and poised enhancers showed reduced H3K27me3 together with increased H3K4me3 and H3K4me1, respectively, compared to CDX2-unbound CREs (**Extended Data Fig. 7f,g**), consistent with remodelling towards a more permissive chromatin state. However, these changes did not correlate with changes in chromatin looping: while loops associated with CDX2-bound poised enhancers were stronger than their CDX2-unbound counterparts, interactions involving bivalent promoters and active enhancers remained comparatively stable and those at active promoters were weaker (**Extended Data Fig. 7h,i**). Thus, changes in CDX2-associated loops were not uniformly coupled to changes in histone modifications.

Collectively, these results suggest that changes in chromatin interactions are only partially coupled to histone modifications at CREs. During posteriorisation, Polycomb-associated interactions are transiently reorganised, while CDX2-associated interactions progressively strengthen across differentiation. However, this global strengthening of CDX2-bound interactions was not uniformly recapitulated at CDX2-bound CREs: despite acquisition of more permissive histone modifications, interaction strength did not vary unidirectionally, suggesting that CDX2 may have overlapping but distinct roles in regulating local chromatin state and chromatin interactions. The emergence of WNT-responsive regulatory elements and CDX2-associated chromatin interactions during this transition further suggests that WNT/β-catenin signalling may regulate chromatin mobility and 3D genome organisation during posteriorisation.

### WNT/ß-catenin signalling rapidly increases chromatin mobility and reorganisation during posteriorisation

To determine whether WNT/β-catenin signalling remodels chromatin before transcriptional responses, we acutely modulated WNT/β-catenin signalling. We added the GSK3 inhibitor CHIRON99021 (CHIRON) to EpiLCs and withdrew it from CEpiLCs before quantifying chromatin mobility, 3D chromatin organisation and transcription over time (**Fig. 3a,b**). This strategy allowed us to distinguish signalling-dependent changes in chromatin organisation from downstream transcriptional responses. Bulk RNA sequencing following CHIRON modulation detected no differentially expressed genes after one hour in either EpiLCs or CEpiLCs (**Extended Data Fig. 8a-c**). Transcriptional responses emerged after six hours, with 104 and 84 differentially expressed genes following CHIRON addition and withdrawal, respectively, including canonical WNT-responsive targets such as *T/Bra*, *Cdx1*, *Cdx2*, *Cdx4*, *Axin2*, *Sp5*, *Sp8* and *Dkk1/4* (**Extended Data Fig. 8d**)^39^. Gene Ontology analysis showed enrichment for WNT signalling, somitogenesis and anterior–posterior axis specification (**Extended Data Fig. 8e**).

**Figure 3.**
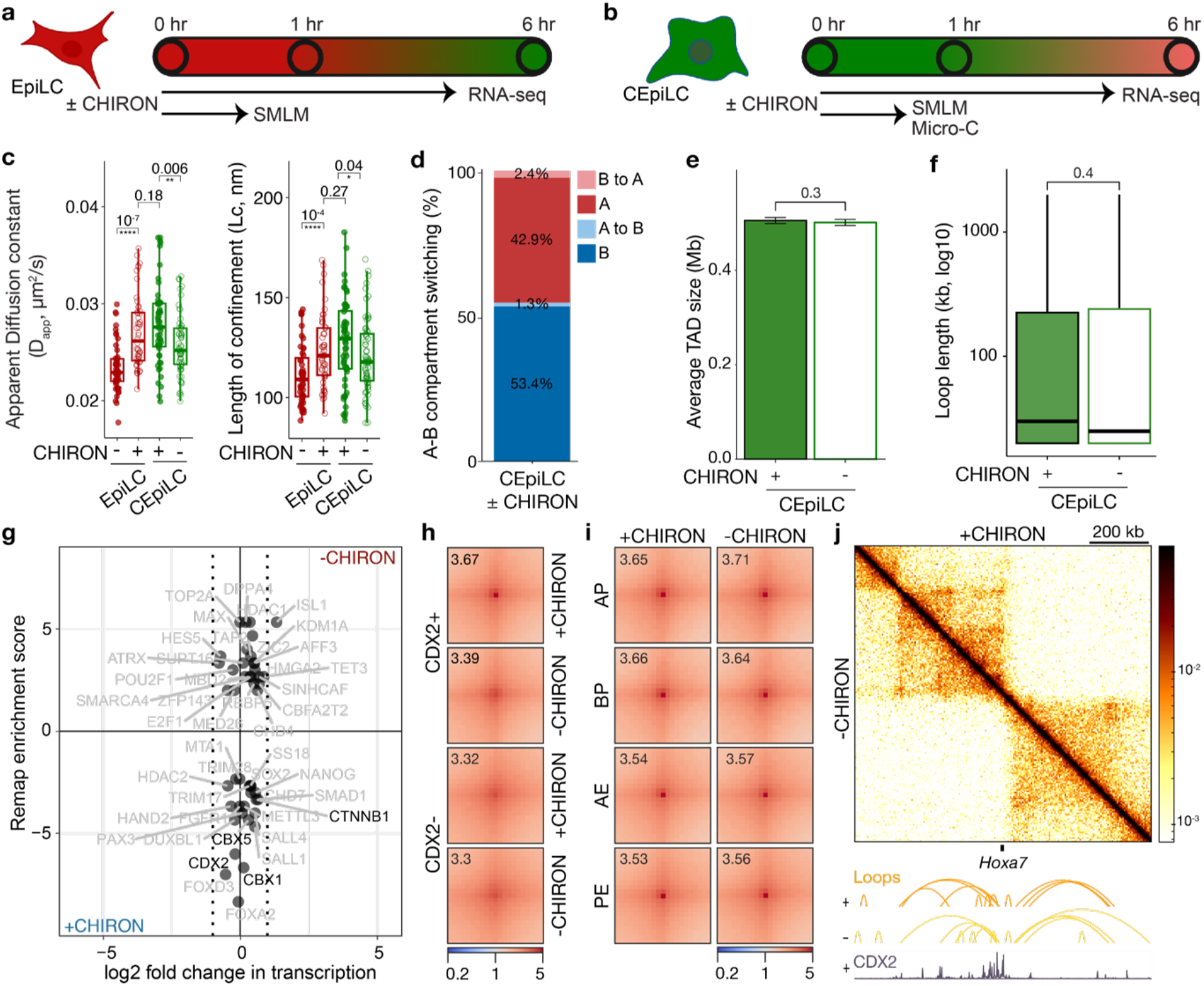
WNT/β-catenin signalling rapidly increases chromatin mobility and reorganisation of 3D enhancer-promoter loops prior to transcriptional changes. **(a-b)** Schematic of CHIRON modulation experiments where CHIRON was both **(a)** added in EpiLCs and **(b)** removed in CEpiLCs, respectively to block WNT/β-catenin signalling. Samples were collected at 0, 1, and 6 hours for SMLM, bulk RNA-seq and Micro-C (CEpiLC only). **(c)** Boxplots quantifying the apparent diffusion constant (D_app_) and confinement radius (Lc) of fast-moving chromatin after modulation of CHIRON in EpiLCs and CEpiLCs. Boxplot fill corresponds to treatment condition with individual datapoints corresponding to trajectories averaged per field-of-view. Box plots show the median (center line), 25th–75th percentiles (box bounds) and 1.5× the IQR (whiskers). Statistical significance was calculated using a Wilcoxon rank-sum test with Holm correction for multiple testing (*n* > 50 FOVs). **(d)** A/B compartment changes after CHIRON removal in CEpiLCs using 1Mb genomic bins. **(e)** Bar plot of TAD size at 600kb resolution in CEpiLCs after CHIRON modulation. Statistical significance was assessed using a two-sided Wilcoxon rank-sum test. **(f)** Box and whisker plot of loop length with and without CHIRON for high-confidence loops identified by FitHiC2 (FDR<10^−5^). Statistical significance was assessed using a two-side Wilcoxon rank-sum test. **(g)** Correlation scatter plot of log2 fold change in transcript levels (bulk RNA-seq) against the ReMap enrichment score (Micro-C) shows changes in enrichment for WNT/β-catenin-associated transcription factors or Polycomb subunits (highlighted in black). The dotted line corresponds to a log2 fold change in transcript levels > 1. **(h)** APA plots +/− CHIRON for CEpiLC loops with CDX2 bound at ≥ 1 anchor in wild-type CEpiLCs (10 kb bins for 100 kb region centred on wild-type CEpiLC loops). **(i)** APA plots stratified by CRE class +/− CHIRON for CEpiLC loops. **(j)** Micro-C contact maps with and without CHIRON at the *Hoxa7* locus. Top, contact maps comparing CEpiLC with CHIRON (top right corner) and without CHIRON (bottom left corner). Bottom, corresponding chromatin loops and CDX2 CUT&RUN tracks for CEpiLC. Scale bar, 200 kb.

**Extended Data Figure 8.**
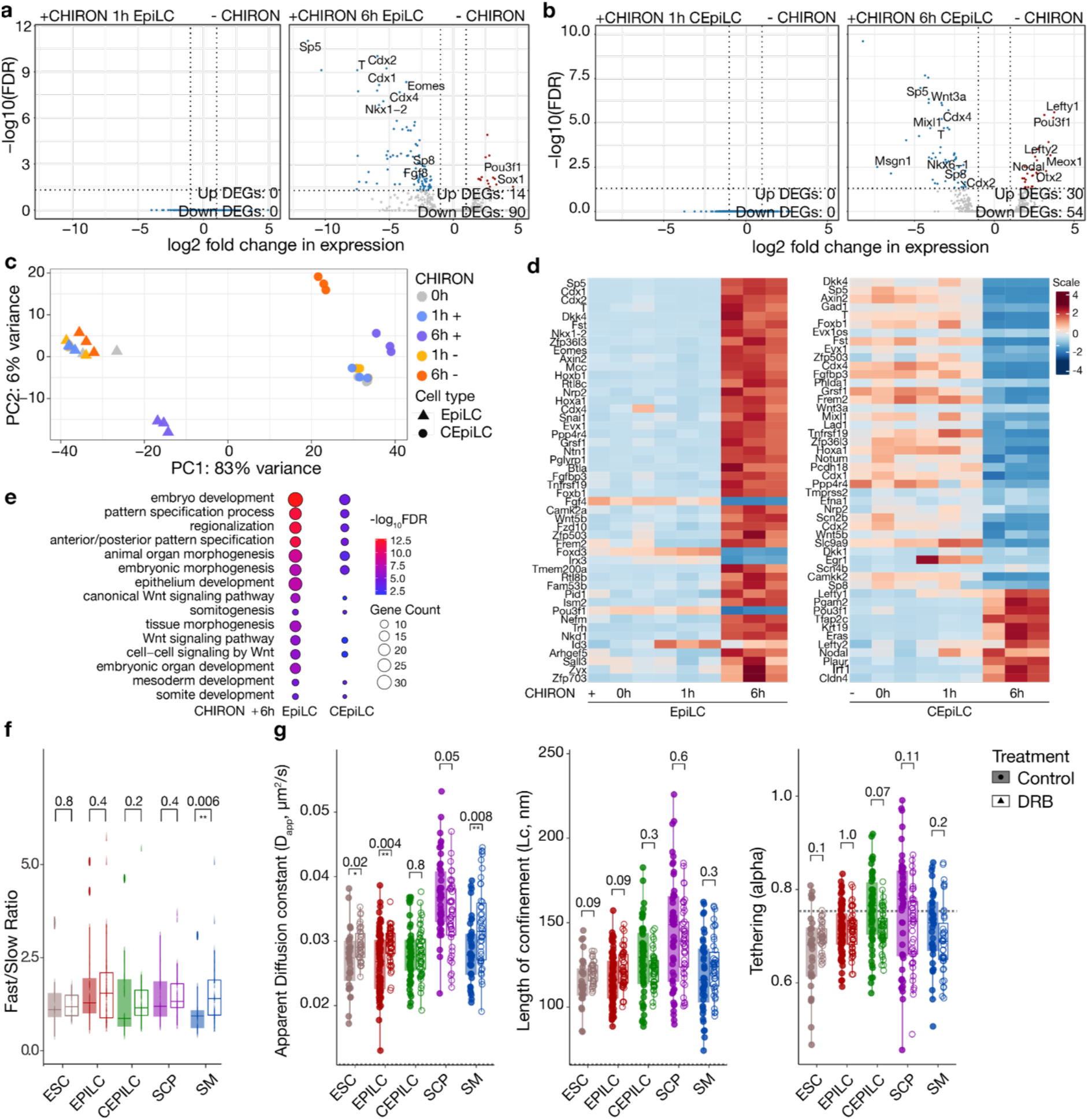
Chromatin mobility does not correlate with transcription. **(a)** Volcano plot of differentially expressed genes (DEGs) in EpiLCs after 1 and 6 hours of CHIRON modulation. **(b)** Volcano plot of DEGs in CEpiLCs after 1 and 6 hours of CHIRON modulation. **(c)** Principal component analysis of RNA-seq samples for EpiLCs following 1 and 6 hr of CHIRON addition and CEpiLCs following 1 and 6 hr of CHIRON removal validate reproducibility and show no transcriptional changes after 1 hr of CHIRON addition/removal after 1 hr of CHIRON addition/removal. **(d)** Heatmap of top 50 differentially expressed genes (DEGs) for (left) EpiLCs following 1 and 6 hr of CHIRON addition and (right) CEpiLCs following 1 and 6 hr of CHIRON removal. Each column represents a biological replicate, and rows represent individual genes. Expression values are mean-centred and scaled per gene (red = upregulated, blue = downregulated). **(e)** Gene Ontology reveals enrichment for WNT-responsive and CEpiLC-associated terms after 6 hours of CHIRON addition in EpiLCs. **(f)** Boxplot showing ratio of fast- vs slow-moving chromatin in WT vs DRB samples for each cell type. Statistical significance tested using two-sided Wilcoxon rank-sum tests followed by Benjamini–Hochberg correction for multiple testing. **(g)** Boxplots show the FOV-averaged apparent diffusion constant (D_app_), confinement radius (Lc) and anomalous exponent (tethering, α) of fast diffusing chromatin trajectories. Colours indicate different cell types. Boxplot fill corresponds to treatment condition. Wilcoxon rank-sum test with Holm correction for multiple testing.

**Extended Data Figure 9.**
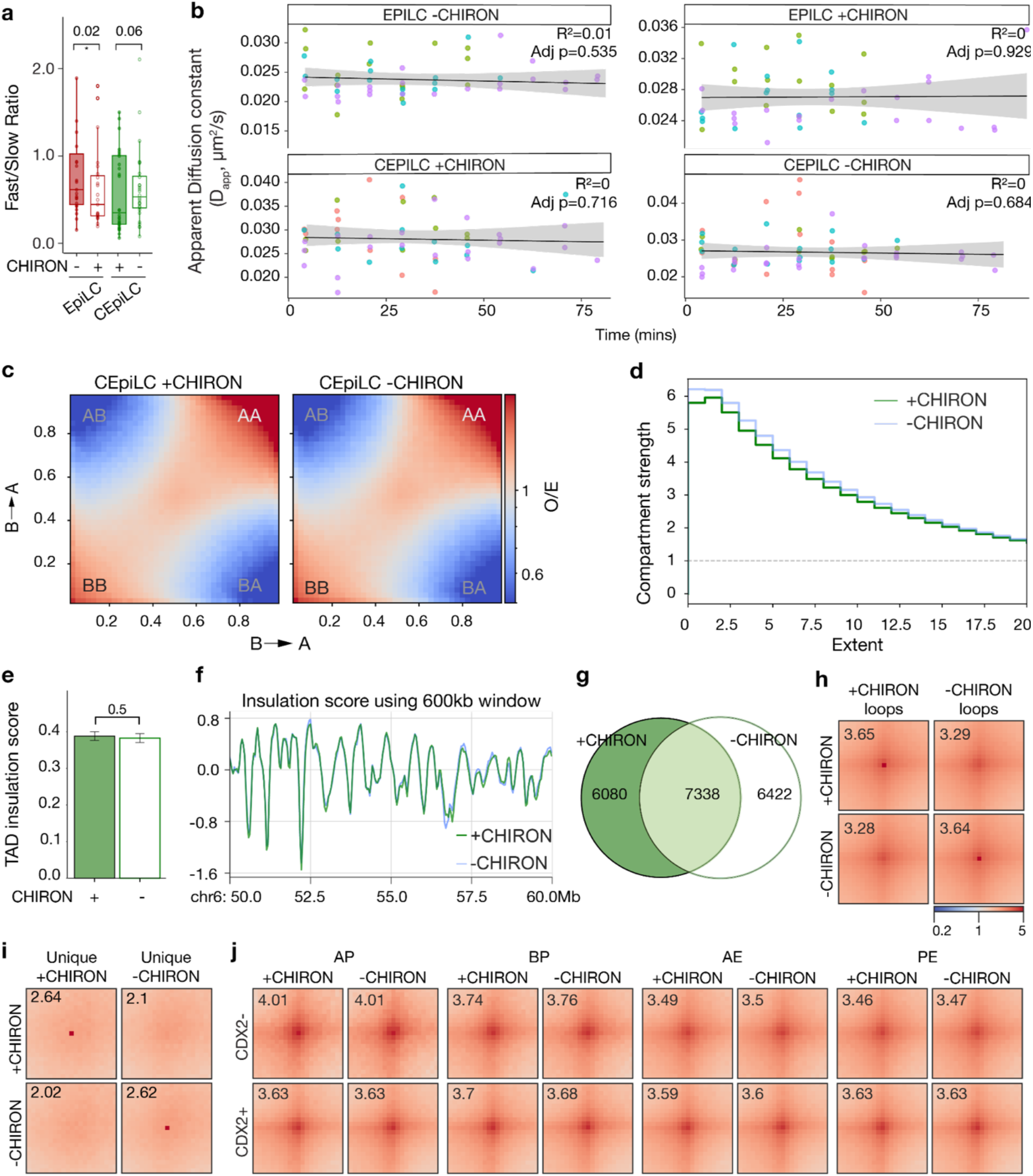
WNT/β-catenin signalling increases chromatin mobility, which correlates with local changes in 3D chromatin looping but no detectable changes in genome compartmentalisation. **(a)** Boxplot showing ratio of fast- vs slow-moving chromatin upon CHIRON addition or removal in EPiLCs or CEpiLCs respectively. Statistical significance was assessed using two-sided Wilcoxon rank-sum tests followed by Benjamini–Hochberg correction for multiple testing. **(b)** Apparent diffusion constant plotted as a function of imaging time for individual fields of view in EpiLCs and CEpiLCs cultured in the presence or absence of CHIRON. Each point represents one field of view, coloured by biological replicate. Solid lines indicate linear regression fits with 95% confidence intervals shown in grey. No significant relationship between imaging time and D_app_ was observed in any condition (adjusted *p* values shown). **(c)** Genome-wide saddle plots calculated as previously described (**Extended Data Fig. 3b**) from Micro-C contact matrices for CEpiLC showing compartment strength during CHIRON modulation. **(d)** Line plot showing compartment strength at different values of extent/genomic scale (in Mb) after CHIRON removal in CEpiLCs, calculated as the ratio (AA + BB)/(AB + BA) from saddle plots of interactions between A and B compartments (see **panel c**), where higher values indicate stronger compartmentalisation. **(e)** Bar plot of TAD insulation score at 600kb resolution in CEpiLCs after CHIRON modulation. Statistical significance was assessed using a Mann-Whitney-Wilcoxon two-sided test. **(f)** Visual representation of the TAD insulation score at the Hox locus on chromosome 6 at 500kb resolution. **(g)** Euler diagram of unique and shared loops with and without CHIRON withdrawal in CEpiLCs. **(h)** APA plots of all loops with and without CHIRON withdrawal in CEpiLCs. **(i)** APA plots of unique loops with and without CHIRON withdrawal in CEpiLCs. **(j)** APA plots of the CRE classes stratified by CDX2-bound status with and without CHIRON withdrawal in CEpiLCs.

To further validate whether transcription can drive changes in chromatin mobility in our specific cell types, we inhibited transcription elongation with 5,6-dichloro-1-β-D-ribofuranosylbenzimidazole (DRB)^31,46^. In CEpiLCs, DRB treatment had no detectable effect on the proportion of slow- and fast-moving chromatin or on any measured biophysical parameter of chromatin mobility (**Extended Data Fig. 8f**). Although DRB altered chromatin mobility in several other cell states, these effects did not recapitulate the increase in chromatin mobility observed during the EpiLC-to-CEpiLC transition (**Extended Data Fig. 8g**), arguing against transcriptional elongation as the primary driver of increased chromatin mobility during posteriorisation. Together, the absence of any mobility change following inhibition of transcription elongation, and the delayed emergence of bulk transcriptional changes in CEpiLCs suggest that a 1 hr time window after CHIRON modulation would allow us to assess how WNT/β-catenin influence chromatin mobility and organisation in the absence of bulk transcriptional change.

We therefore conducted single-nucleosome tracking following acute CHIRON perturbation and showed no change in the proportion of fast- and slow-moving chromatin (**Extended Data Fig. 9a**). However, CHIRON addition significantly increased chromatin mobility in EpiLCs, whereas CHIRON withdrawal reduced mobility in CEpiLCs (**Fig. 3c**). Changes in D_app_ did not correlate with imaging time (**Extended Data Fig. 9b**), suggesting that that WNT/β-catenin signalling can rapidly modulate chromatin mobility within minutes.

We next asked whether these rapid changes in chromatin mobility were accompanied by 3D genome reorganisation. Micro-C performed one hour after CHIRON withdrawal in CEpiLCs, revealed minimal changes in A/B compartment organisation or TAD organisation (**Fig. 3d,e; Extended Data Fig. 9c-f**). By contrast, high-confidence chromatin loops were extensively remodelled (**Extended Data Fig. 9g-i**), consistent with rapid reorganisation of local chromatin interactions rather than large-scale disruption of genome organisation. Although loop length did not significantly change (**Fig. 3f**), ReMap enrichment analysis revealed that loops unique to CHIRON withdrawal were enriched for Polycomb-associated factors, including DPPA4^70^, whereas loops maintained in the presence of CHIRON were enriched for canonical WNT-associated transcription factors, including β-catenin and CDX2 (**Fig. 3g**). Aggregate peak analysis supported the preferential weakening of CDX2-bound interactions following CHIRON withdrawal (**Fig. 3h**), consistent with WNT/β-catenin signalling maintaining a CDX2-associated chromatin interaction landscape. In contrast, interactions associated with CREs showed minimal changes (**Fig. 3i, Extended Data Fig. 9j**), suggesting that WNT/β-catenin-responsive CDX2-associated contacts occur predominantly outside annotated CREs Representative WNT/β-catenin-responsive loci, including *Hoxa7*, similarly showed minimal reorganisation of local contacts (**Fig. 3j**). Together, these findings suggest that WNT/β-catenin signalling maintains a subset of CDX2-associated chromatin interactions while opposing the re-establishment of Polycomb-associated interactions, independently of histone modifications at CREs.

Collectively, these findings identify increased chromatin mobility and reorganisation of CDX2- and Polycomb-associated interactions as early responses to WNT/β-catenin signalling that precede widespread transcriptional changes during posteriorisation.

### CDX factors increase WNT/β-catenin-dependent chromatin mobility and reorganisation

As changes in chromatin mobility and local interactions preceded widespread transcriptional responses, we reasoned that they were likely mediated by factors already associated with WNT/β-catenin-responsive chromatin. CDX2 was enriched at CEpiLC-specific chromatin loops (**Fig. 2e**), CHIRON-dependent loops (**Fig. 3g**), and occupied lineage-associated CREs (**Extended Data Fig. 7c-e**). Together with its established roles in posterior identity and chromatin accessibility^41,42^, these observations identified CDX factors as candidate mediators of WNT/β-catenin-dependent chromatin mobility. We therefore assessed whether CDX factors mediate WNT/β-catenin-dependent changes in chromatin mobility and organisation.

We quantified single-nucleosome mobility following CHIRON modulation in *Cdx1,2,4*-null cells^71^ expressing H2B-HaloTag, as members of the CDX family can have redundant functions^72,73^. These cells acquire an initial T/Bra- and SOX2-expressing progenitor state in response to WNT activation but subsequently fail to undergo appropriate posterior neural or somitic lineage specification (**Extended Data Fig. 10a-d**), consistent with the established role of CDX factors in posterior fate acquisition^71,74,75^. Whereas chromatin mobility was comparable between wild-type and *Cdx1,2,4*-null EpiLCs, *Cdx1,2,4*-null EpiLCs showed no CHIRON-induced increase in chromatin diffusion (D_app_) or confinement radius (Lc) and *Cdx1/2/4*-null CEpiLCs retained EpiLC-like mobility (**Fig. 4a, Extended Data Fig. 10e**). Thus, CDX factors mediate the WNT/β-catenin-dependent increase in chromatin mobility during posteriorisation.

**Figure 4.**
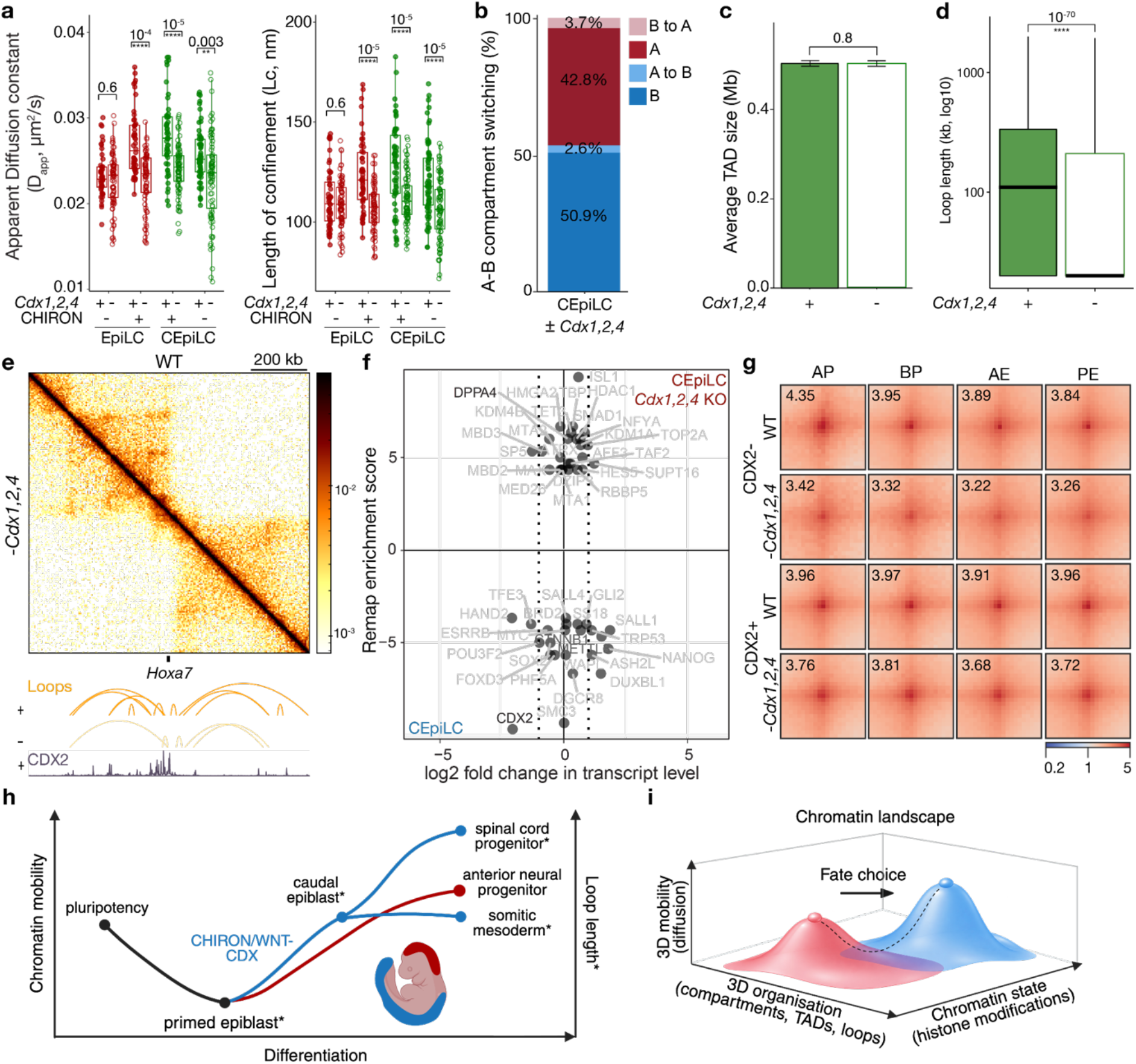
CDX factors mediate increased chromatin mobility and 3D chromatin reorganisation in response to WNT/β-catenin signalling. **(a)** Boxplots showing apparent diffusion constant (D_app_) and confinement radius (Lc) of fast-moving chromatin trajectories for wild-type and *Cdx1,2,4*-null EpiLCs (red) and CEpiLCs (green) upon addition or removal of CHIRON. Dots represent average values per field of view and statistical testing was assessed using Dunn’s post-hoc test with Benjamini-Hochberg correction for multiple testing. **(b)** A/B compartment changes when comparing wild-type to *Cdx1,2,4*-null CEpiLCs using 1Mb genomic bins. **(c)** Bar plot comparing TAD size between wild-type and *Cdx1,2,4*-null CEpiLCs. Statistical testing assessed using a Mann-Whitney-Wilcoxon two-sided test. **(d)** Box and whisker plot of loop length change between wild-type and *Cdx1,2,4*-null CEpiLCs for high-confidence loops identified by FitHiC2 (FDR<10^−5^). **(e)** Micro-C contact maps with and without CDX factors in CEpiLCs at the *Hoxa7* locus. Top, contact matrix comparing WT CEpiLC (top right corner) and *Cdx1,2,4*-null CEpiLCs (bottom left corner). Bottom, corresponding chromatin loops and CDX2 CUT&RUN tracks for CEpiLC. Scale bar, 200 kb. **(f)** Correlation scatter plot of log2 fold change in transcript levels (bulk RNA-seq) against the ReMap enrichment score (Micro-C) when comparing WT and *Cdx1,2,4*-null CEpiLCs shows changes in enrichment for CDX2 or Polycomb-associated proteins (highlighted in black). The dotted line corresponds to a log2 fold change in transcript levels > 1. **(g)** APA plots of loops stratified according to CRE class and CDX2-status. **(h)** Chromatin mobility has similar properties in ESCs, CEpiLCs, SM and ANPs, lowest diffusion in EpiLCs and highest diffusion in SCPs, independent of differentiation status. Diffusion speed correlates with loop length in EpiLCs, CEpiLCs, SM and SCPs (*). CHIRON/WNT modulation drives chromatin mobility changes in EpiLCs and CEpiLCs, and seems to be mediated by CDX factors in CEpiLCs. **(i)** Fate choices are accompanied by changes in the chromatin landscape, including histone marks, 3D chromatin organisation (A/B compartments, TADs, loops), and 3D chromatin mobility.

**Extended Data Figure 10.**
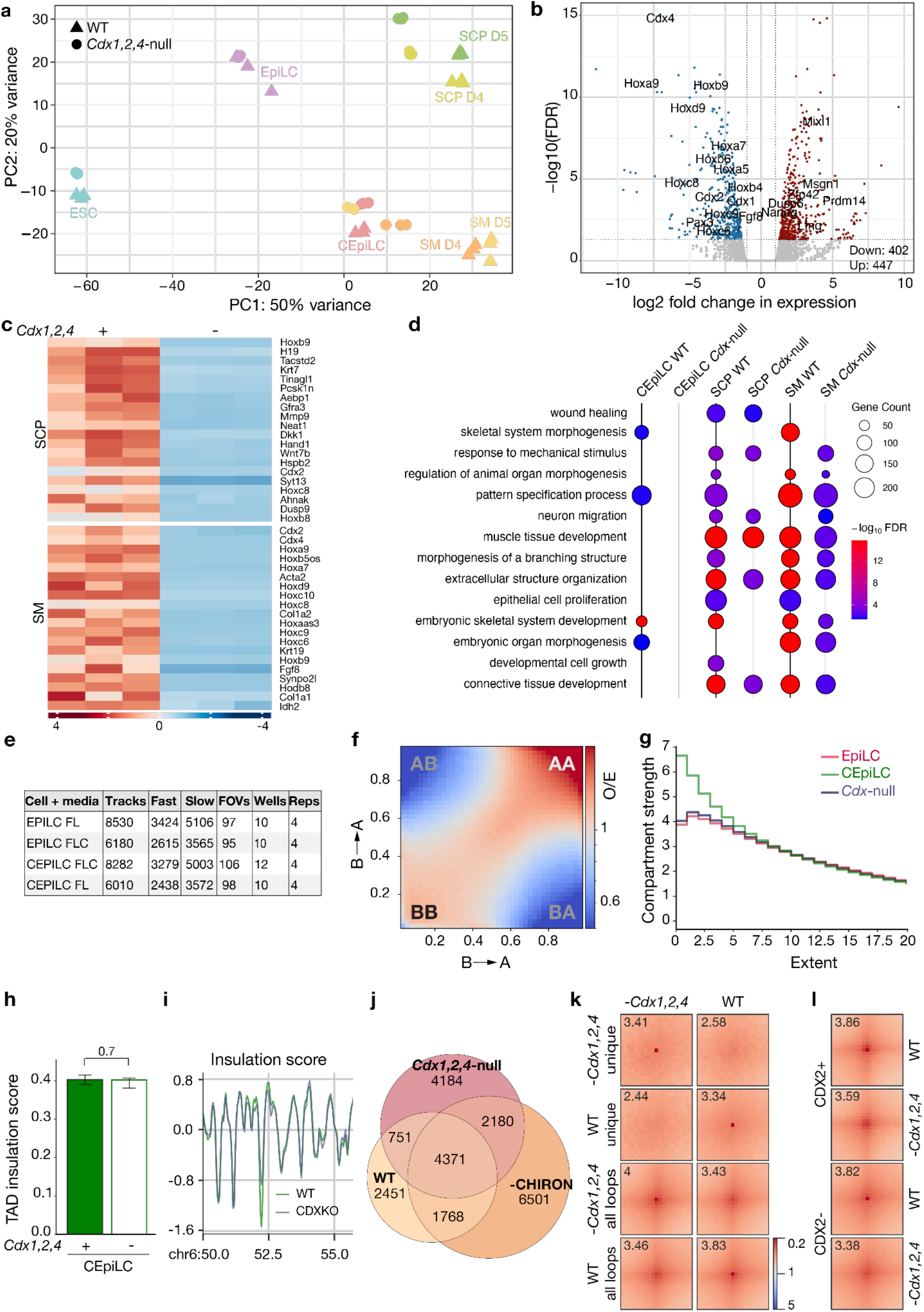
CDX factors are required for 3D genome reorganisation at posterior genes during posteriorisation and differentiation toward SM and SCP fates. **(a)** PCA plot comparing RNA-seq of wild-type and *Cdx1,2,4*-null cells at each stage of differentiation reveals inability to acquire SM fate and large gene expression changes in SCPs. **(b)** Volcano plots of differentially expressed genes (DEGs) when comparing wild-type and *Cdx1,2,4*-null cells grown in conditions that give rise to CEpiLCs, SCPs and SM. **(c)** Heatmap of top DEGs in wild-type and *Cdx1,2,4*-null SM and SCP cells. **(d)** GO term plots based on the DEGs in wild-type and *Cdx1,2,4*-null CEpiLC, SM and SCP cells. **(e)** Table showing number of unclassified (total) and classified (slow/fast) trajectories, total FOVs, wells and biological replicates per dataset. **(f)** Genome-wide saddle plot derived from Micro-C contact matrices for *Cdx1,2,4-*null CEpiLCs. **(g)** Line plot of the compartment score profiles across chromosome 6 (chr6) for *Cdx1,2,4-*null CEpiLCs compared to WT EpiLCs and CEpiLCs. **(h)** TAD insulation score at 600kb resolution in CEpiLCs after CHIRON modulation. Statistical significance assessed using a Mann-Whitney-Wilcoxon two-sided test. **(i)** Line plot of the TAD insulation score calculated for *Cdx1,2,4-null* and WT CEpiLCs with pairwise comparison highlighting genomic regions undergoing TAD changes at the chromosome 6 *Hox* region. **(j)** Euler diagram showing the overlap between significant chromatin loops detected during multiway comparisons between WT, CHIRON removal and *Cdx1,2,4-*null CEpiLCs. **(k)** APA plots of all and unique loops in WT and *Cdx1,2,4*-null CEpiLCs. **(l)** APA plots of CDX2-bound and independent loops in WT and *Cdx1,2,4*-null CEpiLCs.

Micro-C of *Cdx1,2,4*-null CEpiLCs revealed minimal A/B compartment switching and little change in TAD organisation relative to wild-type CEpiLCs (**Fig. 4b,c; Extended Data Fig. 10f-i**) but lacked the increased B-B interactions observed during posteriorisation (**Extended Data Fig. 3b**). In contrast, chromatin loops were extensively remodelled and significantly shorter in *Cdx1,2,4*-null CEpiLCs (**Fig. 4d,e; Extended Data Fig. 10j,k**). ReMap analysis showed that loops lost in *Cdx1,2,4*-null cells were enriched for CDX2, SMC3 and additional lineage-associated transcription factors, whereas gained loops were enriched for Polycomb-associated regulators, including DPPA4 (**Fig. 4f**), resembling both EpiLCs (**Fig. 2e**) and CEpiLCs following CHIRON withdrawal (**Fig. 3g**). APA of loops stratified by CRE class and wild-type CDX2 occupancy revealed that loops associated with CREs were overall stronger in *Cdx1,2,4*-null cells than in wild-type, irrespective of whether they were CDX2-bound or CDX2-unbound (**Fig. 4g; Extended Data Fig. 10l**), suggesting that CDX transcription factors have both direct and indirect roles in maintaining loops at CREs. Together, these results identify CDX transcription factors as mediators of WNT-dependent changes in chromatin mobility and lineage-associated chromatin interactions during posteriorisation.

## Discussion

Our results identify developmental signalling as a rapid regulator of chromatin mobility and place this physical transition upstream of widespread transcriptional responses during posteriorisation. Contrary to prevailing models in which chromatin mobility progressively decreases during differentiation^34,38^, chromatin diffusion was lowest in EpiLCs and highest in posterior SCPs, varying in a cell type- and anterior–posterior-specific manner. Together, these findings establish that chromatin mobility does not simply decline as pluripotent cells differentiate but is instead dynamically regulated according to lineage and positional identity **(Fig. 4h)**.

Chromatin mobility determines how the chromatin polymer explores its conformational landscape^76–79^ and so changes in mobility alter the repertoire of chromatin conformations sampled by genomic loci over time, and more broadly, the energetic landscape explored by the chromatin fibre^80–82^. A recent study by Mazocca et al (2026) that compares chromatin mobility across cell lines provides important context for interpreting our observations^83^. It supports our finding of cell type-specific differences in chromatin mobility but also highlights a fundamental constraint: while highly subdiffusive chromatin mobility enables remarkably efficient diffusive search over local distances (<200-300 nm), target search over larger distances rapidly becomes inefficient and is likely to require facilitation. This at first appears difficult to reconcile with the large-scale 3D genome reorganisation known to occur during development^7,84,85^. However, our observation that WNT/β-catenin signalling and CDX factors increase chromatin mobility provides a framework for how extensive 3D genome reorganisation may be facilitated during developmental cell state transitions, consistent with recent work showing that CDX2 overexpression increases chromatin mobility^35^. Rather than directly specifying regulatory interactions, the WNT-CDX-dependent increase in chromatin mobility during posteriorisation likely facilitates the exploration of alternative chromatin conformations, increasing opportunities for new interactions between genomic regions, including cis-regulatory elements. Cell type-specific chromatin interactions can subsequently emerge through lineage-specific chromatin regulators progressively stabilising a subset of these conformations. Nevertheless, the WNT-CDX axis identified here is unlikely to be the sole determinant of chromatin mobility, as mobility increased further during SCP specification. Distinct developmental contexts therefore likely regulate chromatin mobility through different regulatory mechanisms.

By comparing imaging and Micro-C analyses across the same cell states, we reveal that chromatin mobility and 3D genome organisation are correlated but distinct features of cell state. Differentiation was accompanied by coordinated changes in chromatin mobility and loop length, consistent with our previous work showing that chromatin regulators such as NuRD and MLL2 can increase or decrease chromatin mobility to facilitate the distal or local exploration of cis-regulatory elements respectively^44,46^. In contrast, chromatin mobility correlated poorly with genome compartmentalisation, continuing to increase during SCP specification despite comparatively modest changes in A/B compartments and TADs. Moreover, histone modifications and chromatin loop reorganisation were uncoupled across multiple state transitions, indicating that these layers of genome organisation are regulated independently. Chromatin mobility, histone modifications and 3D interactions therefore represent distinct but interacting regulatory properties of the chromatin during a cell state transition **(Fig. 4i)**.

The ability of developmental signals to modulate chromatin mobility may have broader implications for development and disease. Increased mobility may enhance interactions of regulatory elements, but greater conformational exploration could also increase inappropriate enhancer–promoter communication or susceptibility to genome instability. Dysregulation of both WNT/β-catenin signalling and CDX factors is common in cancer^86,87^, and transient loss of Polycomb function has recently been implicated in malignant transformation^88^, raising an intriguing parallel with the transient reduction in Polycomb-associated interactions observed during posteriorisation. Whether similar mechanisms are co-opted during disease remains an important question for future studies. Together, our findings establish chromatin mobility as a signalling-responsive physical property of cell state and suggest that modulation of the conformational landscape explored by the chromatin fibre provides an additional biophysical layer through which extracellular signals shape genome organisation during development.

## Methods

**Plasmids used:**

| Plasmid Name | Description | Source |
| --- | --- | --- |
| LZ10 PBREBAC-H2BHaloTag | Plasmid for stable random integration of H2B-HaloTag sequence using the PiggyBac transposon system. Includes an ampicillin and a neomycin resistance cassette. | Addgene #91564<sup>89</sup> |
| pCyL43 (PBase) | Plasmid expressing PBase, the PiggyBac transposase. Includes an ampicillin resistance cassette. | <sup>90</sup> |

**Mouse embryonic stem cell (mESC) lines:**

| Name in manuscript | Description | Source |
| --- | --- | --- |
| ES-E14tg2a | wild-type ES cell-line derived from a male mouse strain 12910la | <sup>91</sup> |
| ES-E14:H2B-HaloTag | ES cell-line derived from a male mouse strain 12910la stably expressing randomly integrated HaloTag-tagged H2B | This paper |
| *Cdx*<sup>1,2,4</sup>-null | *Cdx1/2/4* triple knock-out (Gouti et al. 2017) | Kindly gifted by Prof. Briscoe |
| *Cdx*<sup>1,2,4</sup>-null:H2B-HaloTag | ES cell-line with *Cdx1,2,4* triple knock-out stably expressing randomly integrated HaloTag-tagged H2B | This paper |

**Sequencing datasets:**

Micro-C (this study)

RNA-seq (this study)

CUT&RUN (this study)

CUT&Tag (this study)

**Software:**

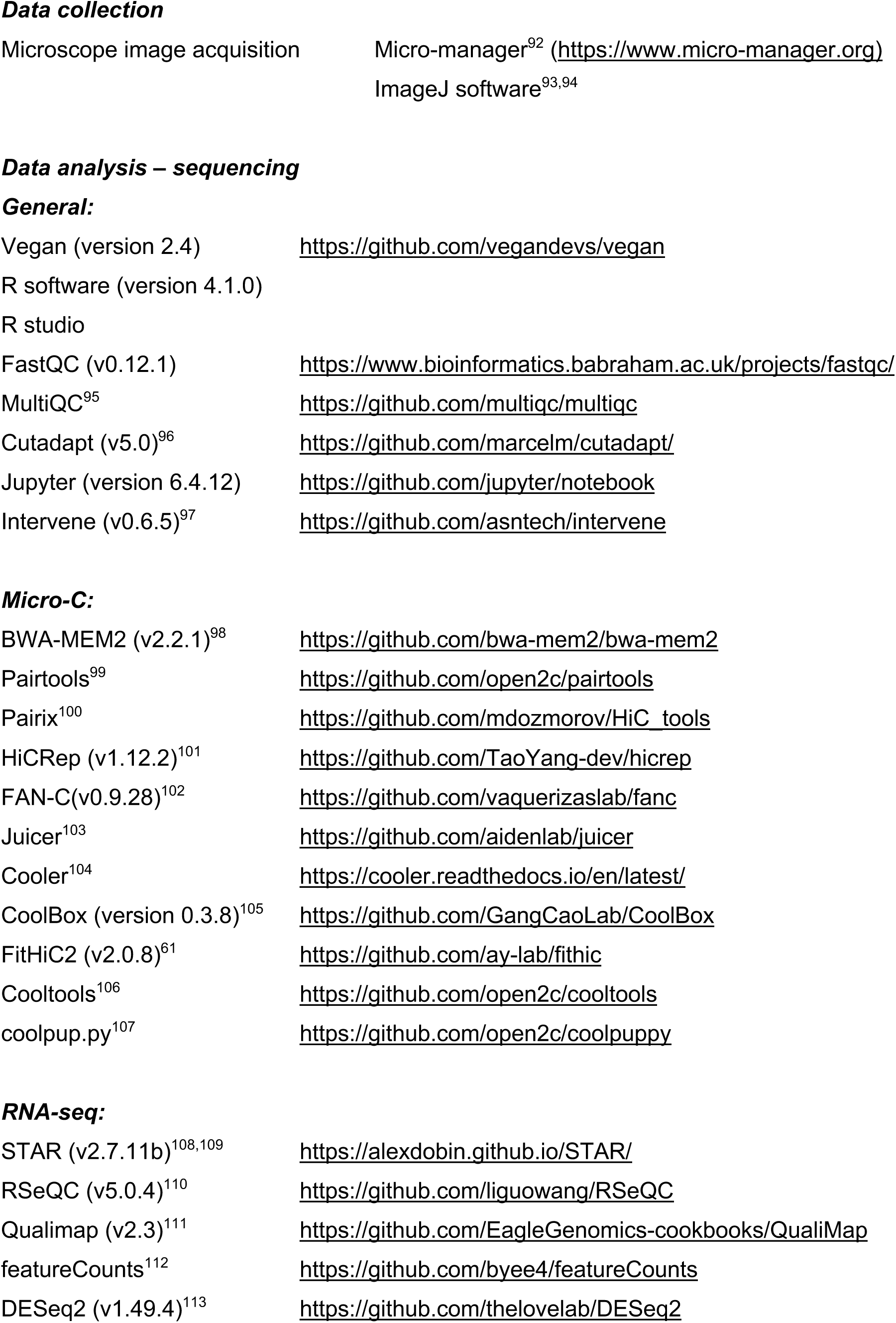
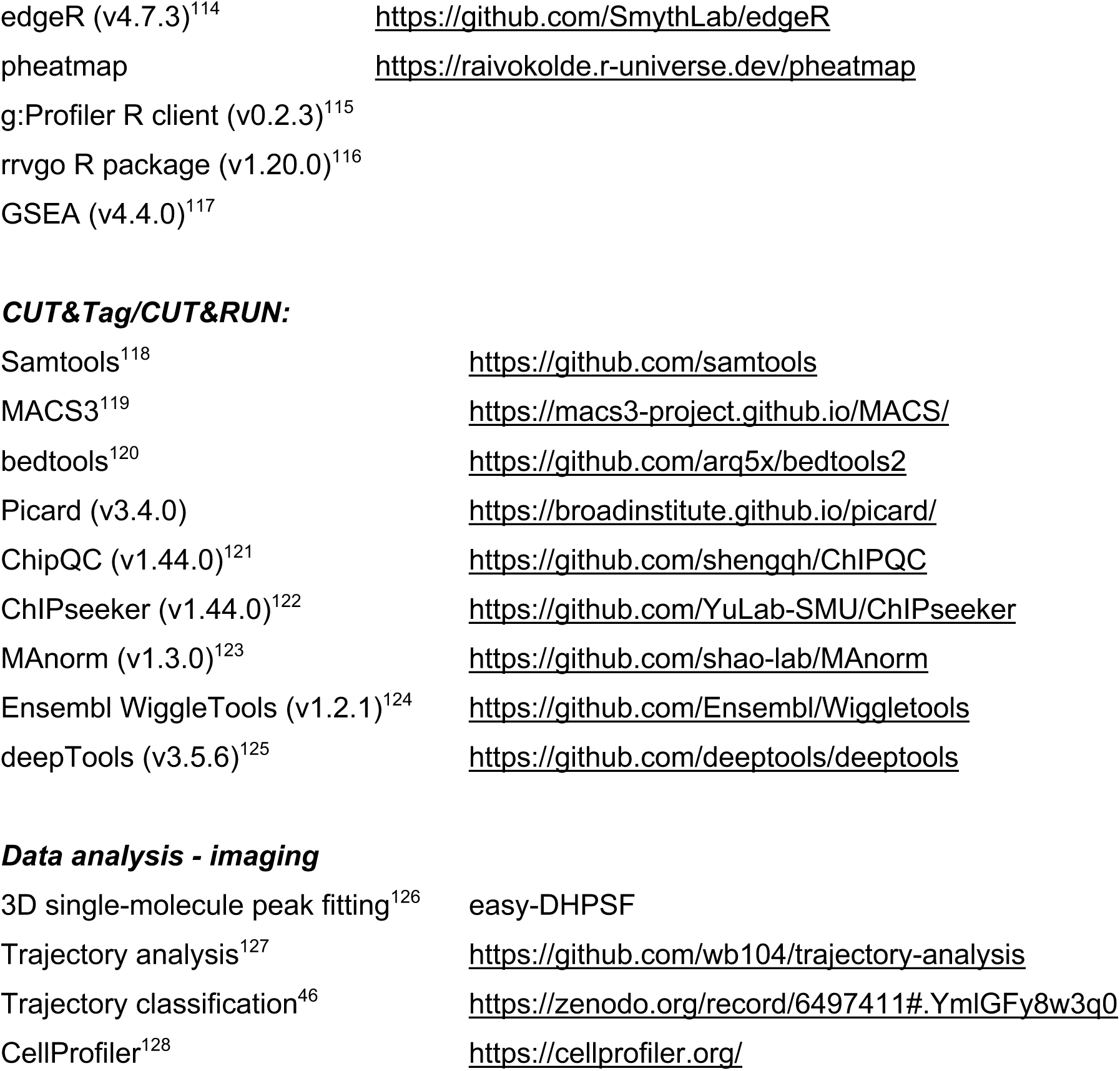

### 1. Tissue culture

#### Embryonic stem cell (ESC) culture

ESCs were grown in phenol red-free 2iLIF medium at 37 °C, 5 % CO_2_ and 80% humidity as previously described^44,129^. All cell lines were routinely screened for mycoplasma contamination and tested negative.

#### Differentiation

ESCs were differentiated to epiblast-like cells (EpiLCs), then caudal epiblast-like cells (CEpiLCs), and finally somitic mesoderm and spinal cord progenitor cells *in vitro* as previously described^42^. Briefly, cells were plated at a density of 5×10^3^ cells/cm^2^ on 0.1% gelatin-coated CellBIND plates (Corning #3335) and cultured in N2B27 supplemented with 10 ng/ml FGF2 and 5 µM LGK974 (Sigma Aldrich 5310910001) (FL media) for 48 hours to EpiLCs. N2B27 was then supplemented with 10 ng/ml FGF2, 5 µM LGK974, and 5 μM CHIR99021 (FLC media) for an additional 24 hours to produce CEpiLCs. Somitic mesoderm and posterior spinal cord progenitors were differentiated by culturing cells for an additional 48 hours in N2B27 supplemented with 5 µM LGK974 and 5 μM CHIR99021 (LC media), or N2B27 with 5 µM LGK974 (L media), respectively, with media changes every 24 hr. Anterior neural cells were generated as previously described with minor changes^41^. Briefly, cells were plated as above up to EpiLC-stage, then at 48 hr the media was changed to FL media, and at 72 hr, changed to N2B27 supplemented with Smooth Agonist (SAG, Cambridge Bioscience #CAY11914) for an additional 24 hr prior to SMLM. Cells were cultured at 37 °C and 5 % CO_2_.

For microscopy experiments, ESCs were plated on 20 μg/mL fibronectin-coated 35 mm Poly-D-Lysine Coated Coverslips (MatTek Corporation P35GC-1.5-14-C). All other conditions were kept the same. For SMLM experiments, somitic mesoderm cells, spinal cord progenitors and anterior neural progenitors were imaged on day 4 as higher cell density on day 5 caused excessive background signal.

#### Cell line generation

Wild-type and *Cdx1,2,4* knockout ESCs expressing HaloTag-tagged histone H2B were generated by transfecting cells with the LZ10 PBREBAC-H2BHalo (Addgene, Plasmid #91564)(Li et al., 2016) and PiggyBAC transposase PBase plasmids (kindly gifted by Dr Brian Hendrich)(Wang et al., 2008) as previously described^44^. 1 µg total DNA of the purified plasmids was added to freshly passaged ESCs using Lipofectamine-3000 (Thermo Fisher, #L3000001) following manufacturer recommendations. Cells were passaged after 24 hours onto a T150 plate in the presence of geneticin (G418, 400 μg/ml, in-house) for 48 hours. G418-resistant cells were then collected for further selection and validation. Antibiotic-resistant cells were labelled with JFX_549_ (Janelia Fluor, 500 nM) for 15 min, washed with PBS, incubated for 30 min in fresh media, then washed, detached with Accutase, and single cell dissociated for FACS. Cells were sorted on a BD FACS Aria™ Fusion Flow Cytometer by the JCBC FACS facility. Positive single clones were collected in 2iLIF + 10% Pan-Serum (in-house) in a 96-well plate for clonal expansion. Clones were then further assayed to ensure pluripotency and differentiation by RT-qPCR and immunofluorescence, respectively.

### 2. Assays

#### Immunofluorescence

For immunofluorescence, cells were differentiated on fibronectin-coated 35 mm Poly-D-Lysine Coated Coverslips (MatTek Corporation P35GC-1.5-14-C) as previously described. Cells were fixed in 4% paraformaldehyde (PFA) for 10 minutes at RT, then permeabilised and blocked with 3% donkey serum in 0.1% Triton/PBS (PBST) for 2 hours at RT. Cells were incubated with primary antibodies in PBST overnight at 4°C. Cells were washed three times 10 minutes with PBST prior to a 2-hour incubation with secondary antibodies in PBST at RT. Cells were washed three times 10 minutes in PBST. Cells were finally incubated with 1:1000 DAPI (Sigma-Aldrich) 10 minutes at RT. All cells were stored at 4°C in fresh PBS 24 hours prior to imaging. Cells were imaged on a LSM980-Airyscan2 (CSCI imaging facility) or LSM710 (CSCI imaging facility).

**Primary Antibodies for immunofluorescence:**

| Target | Species | Concentration | Manufacturer | Catalogue Number | Lot # |
| --- | --- | --- | --- | --- | --- |
| SOX1 | Rabbit | 1:100 | Cell Signalling Technologies | 4194S | 2 |
| SOX2<br>(Btjce) | Rat | 1:200 | Thermo Scientific | 14-9811-82 | 2781641 |
| T/BRA | Goat | 1:100 | Novus Biologicals | AF2085 | KQP0719121 |
| TBX6 | Rabbit | 1:100 | Abcam | AB38883 | 1065600-1 |
| CDX2 | Rabbit | 1:200 | Abcam | AB76541 | GR3380169-17;<br>1000616-51 |

**Secondary Antibodies for immunofluorescence:**

| Fluorophore | Dilution | Host species | Reactivity | Manufacturer | Cat # | LOT |
| --- | --- | --- | --- | --- | --- | --- |
| DyLight488 | 1:500 | Donkey | anti-goat | DyLight | SA5-10086 | WE3275103 |
| CF568 | 1:500 | Donkey | anti-rat | Biotium | 20092 | 16C0912 |
| CF647 | 1:500 | Donkey | anti-rabbit | Biotium | 20047 | 17C0123 |

#### Single Molecule Localisation Microscopy (SMLM)

<u>C</u>ells were differentiated on fibronectin-coated 35 mm Poly-D-Lysine Coated Coverslips (MatTek Corporation P35GC-1.5-14-C). On the day of imaging, H2B-Halo expressing cells were labelled with 50 nM PA-JF_549_ Halo ligand dyes as previously described (dyes were a kind gift from Luke Lavis, Janelia Research Campus)^43,44^. Transcription elongation was inhibited by adding 100 μM 5,6-dichloro-1-β-d-ribofuranosylbenzimidazole (DRB) 2 hr before imaging as previously described^46^. CHIRON modulation was achieved by changing the media to FL or FLC in EpiLCs and CEpiLCs at the start of imaging.

Samples were imaged using a highly inclined and laminated optical sheet (HILO) illumination and double-helix point spread function (DHPSF) microscopy^45,46^. The excitation beams were circularly polarised, collimated and focused to the back plane of an index-matched 1.2 NA water immersion objective lens (Plan Apo VC 60×, Nikon, Tokyo, Japan). A double-helix phase mask optimised for 580 nm (DoubleHelix, Boulder, CO, USA) was positioned in a Fourier plane of the emission path of a fluorescence microscope (Eclipse Ti-U, Nikon) to generate the DHPSF. The objective lens was mounted onto a high-precision motorised linear XY stage (HLD117NN, Prior) with a scanning piezo stage (P726 PIFOC, PI, Karlsruhe, Germany) to control movement in the *xyz* planes. A 4*f* system of lenses relayed the image onto an EMCCD camera (Evolve Delta 512, Photometrics, Tucson, AZ). The effective pixel size is 278 nm. Excitation illumination was provided by a 561 nm laser (561 nm: 200 mW, 06-DPL, Cobolt). Fluorescence signal was separated from the excitation beams into the emission path by a quad-band dichroic mirror (Di01-R405/488/561/635-25×36, Semrock) before being focused into the image plane by a tube lens (TTL200-A, Thorlabs). Long-pass and band-pass emission filters (BLP02-561R-25 and FF01-580/14-25, Semrock) were placed before the camera. Electron multiplication gain was 250. Axial calibration was achieved by imaging a z-stack of fluorescence microspheres (Thermo Fisher Scientific, #T7279) in PBS on a glass slide every 100 nm over a 4µm z-range, 30 ms exposure, 10 images per z position. Using continuous 561nm excitation, fluorescence images were collected of the PA-JF_549_ fluorophores at 500 ms exposure with a power density of ∼160 W/cm^2^. 800-1000 frames were acquired per field of view, with 5-10 FOV per sample, 3 samples per condition, per biological replicate. Every experiment was repeated with at least 3 biological replicates.

#### Micro-C

Micro-C was performed using the Dovetail Micro-C Kit (Dovetail Genomics, #21006) following the manufacturer’s instructions with slight modifications, as described below. Briefly, 1 million cells were harvested, washed in 1x PBS, pelleted and the pellets placed at −80 °C for at least 30 min before proceeding with the protocol. Long-range chromatin crosslinking was achieved by addition of DSG (disuccinimidyl glutarate; ThermoFisher Scientific #A35392) for 10 minutes at RT on a rotator (20 RPM), followed by 37 % formaldehyde (Sigma-Aldrich, #F8775) for 10 minutes at RT to stabilise shorter-range DNA-protein and protein-protein interactions. After two wash steps, cells were digested *in situ* with 2 uL micrococcal nuclease (MNase). EGTA was added to terminate the reaction after exactly 15 min. Cells were subsequently lysed using SDS and the lysate stored at −20 °C for further processing. The size distribution of the MNase-digested DNA was measured to ensure 40-70 % of DNA was between 50-250 bps as recommended and that there was no variation between samples in MNase accessibility. Briefly, a small sample taken from the lysate was subjected to crosslink reversal and Proteinase K digestion to remove protein contaminants. DNA was purified using the Zymo Research DNA Clean and Concentrator-5 Kit (Zymo Research, #D4013) and analysed using an Agilent D5000 ScreenTape (Agilent, #5067-5588) on the Agilent TapeStation 4200 (JCBC NGS facility; Analysis Software Version 5.1). DNA concentration was determined using a Qubit 2.0 Fluorometer (JCBC NGS facility; Qubit^TM^, ThermoFisher Scientific). For proximity ligation, 1000 ng of each lysate was thawed on ice and immobilised on chromatin capture beads. Following two washes, an end polishing step was performed to fill ‘sticky ends’ generated by MNase digestion. After another wash, crosslinked fragments were ligated in a two-step reaction using the provided bridge, bridge ligase and the intra-aggregate ligation reagents. Consequently, a combined crosslink reversal and Proteinase K digestion step was carried out and resulting ligated fragments were eluted from the beads. This step also introduced a biotin-label into the ligated fragments, crucial for enrichment of ligated fragments using streptavidin beads, as described below. DNA was subsequently purified using SPRIselect beads (Beckman Coulter # B23319) at a 1.8-fold bead-to-sample ratio to ensure maximum recovery of all DNA fragments ≥100 bp and stored overnight at −20 °C. Purified and proximity-ligated fragments were subjected to End Repair, followed by Ligation with Illumina Adapters, USER (uracil-specific excision reagent) digest, and another SPRIselect bead-based purification (2.2x bead-to-sample ratio). Ligated sample-adapter fragments were then immobilised on streptavidin beads (via the biotinylated bridge introduced before) and washed for the indicated times with several provided buffers (LWB, NWB, 1x Wash Buffer). Samples were indexed using a set of eight unique UDI Primer Pairs in a standard PCR reaction (12 cycles) as recommended. Primer Pairs were allocated as follows:

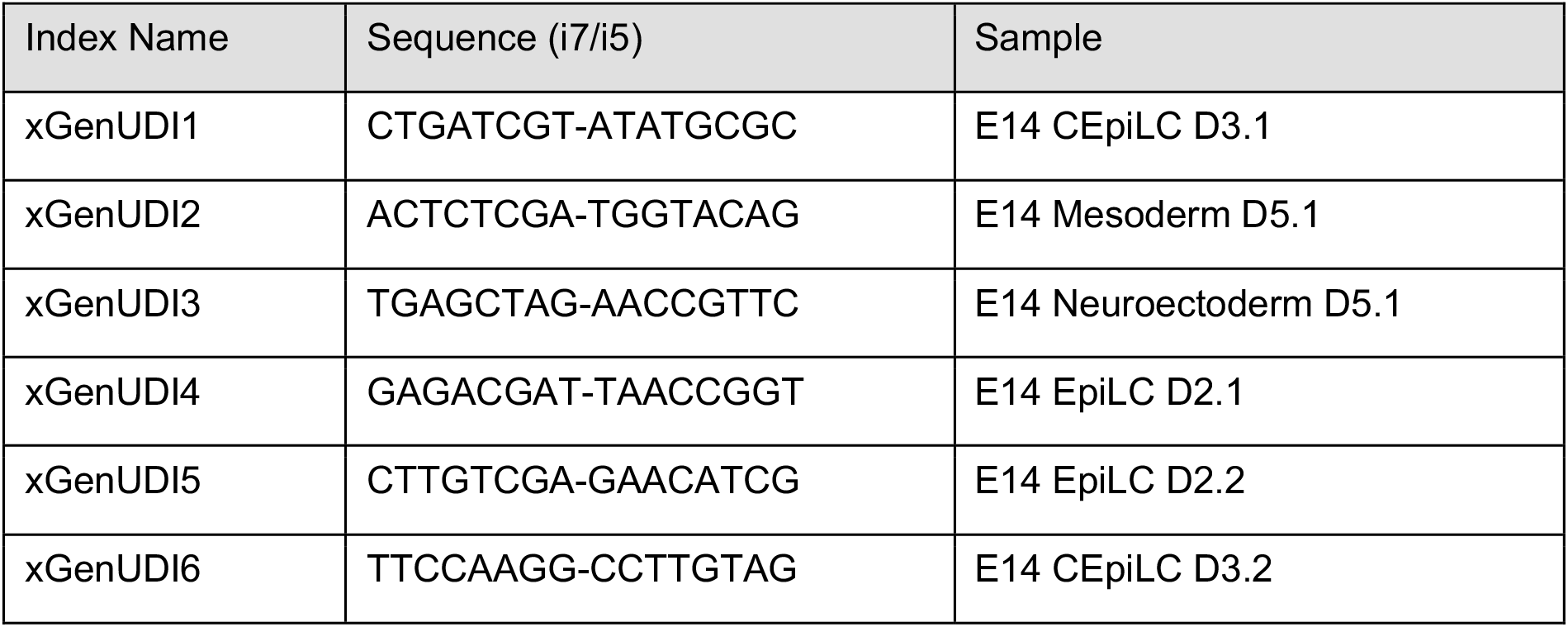
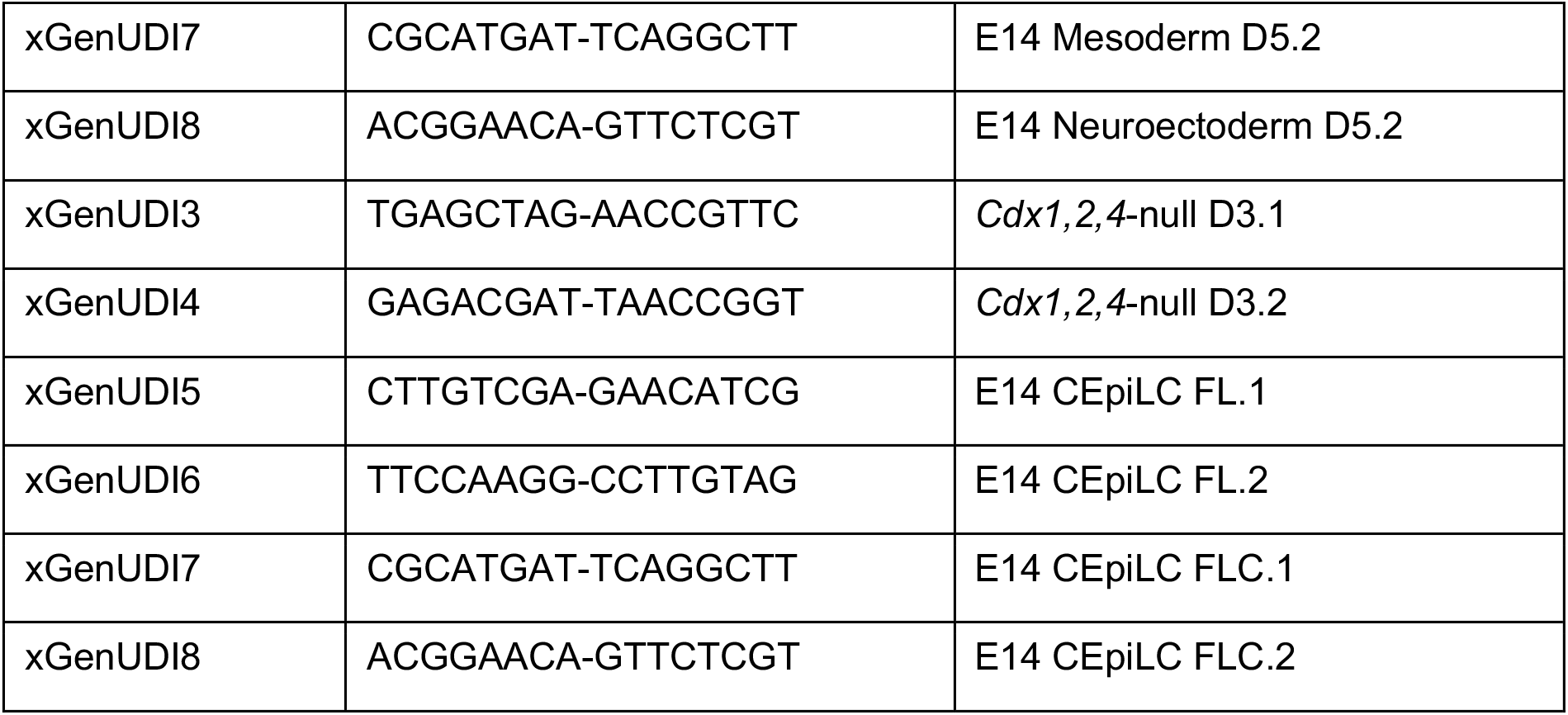

The resulting libraries were subjected to a double-sided selection using SPRIselect beads (right-sided selection at 0.5x, followed by left-sided selection of supernatant at 0.7x bead-to-sample ratio), which excludes any fragments sized <350 bp or >1000 bp. A small sample of each library was run on an Agilent TapeStation 4200 to validate that we had the required fragment size distribution and a Qubit 2.0 Fluorometer to determine DNA concentration. Eight libraries (corresponding to UDI primer pairs 1-8) were pooled at equal concentration to yield a final concentration of 10 nM and the average fragment size of the pooled library was determined using the Agilent TapeStation 4200. Samples were submitted for 150 bp paired-end sequencing at the CRUK-CI Genomics Core Sequencing Facility and sequenced on a NovaSeq 6000 S4 Flow Cell using a standard Illumina sequencing workflow.

| Library Name | Library Barcode | Samples | Average Length [bp] |
| --- | --- | --- | --- |
| E14-pooled | SLX-22776 | E14 EpiLC 1&2, CEpiLC 1&2, Mesoderm 1&2, Neuroectoderm 1&2 | 447 |
| *Cdx1,2,4*-null-pooled | SLX-24169 | *Cdx1,2,4*-null CEpiLC 1&2, E14 CEpiLC FL 1&2, CEpiLC FLC 1&2 | 480 |

#### CUT&Tag

Cleavage Under Targets & Tagmentation (CUT&Tag) was performed using a modified version of the published protocol^68^. Briefly, cells were differentiated on CellBIND dishes as previously described, and 100.000 cells were collected per antibody replicate, in duplicate per condition. Cells were resuspended in Wash Buffer to a concentration of 1 million cells per ml. Cells were subsequently frozen in 10% DMSO for further use. Thawed cells were bound to 1:10 volume BioMag Plus Concanavalin A beads (Cliniscience Ltd #86057-10) previously activated with Binding Buffer for 10 minutes at RT in DNA lo-bind Eppendorf tubes. Cells were subsequently incubated with primary antibodies in Antibody buffer overnight at 4°C with gentle rocking, then secondary antibody in 0.05% Digitonin-Wash Buffer for 1 hour at 4°C with gentle rocking. Samples were then mixed with 1:100 pAG-Tn5 pre-loaded adapter complex in Dig300-wash buffer for 1 hour at 4°C with gentle rocking. The transposase was activated by incubation in Tagmentation Buffer containing magnesium for 1 hour at 37°C. Beads were resuspended in 1% SDS-Proteinase K Release solution and incubated at 37 °C for 1 hour. DNA fragments were cleaned up using 2.2x SPRIselect beads (Beckman Coulter # B23319). 21 mL of eluted DNA was mixed with 2 µL each of 10 mM unique i5 and i7 barcodes, followed by 25 µL NEBNext HI-FI PCR mix, and the mix was amplified by PCR for 12 cycles. After the PCR, the DNA was cleaned-up using 1.1x SPRIselect beads (Beckman Coulter # B23319) and resuspended in 10 mM Tris-HCl pH 8 for quality control using an Agilent D5000 ScreenTape (Agilent, #5067-5588) on the Agilent TapeStation 4200 (JCBC NGS facility; Analysis Software Version 5.1) and a Qubit 2.0 Fluorometer. Pooled samples were submitted for 150 bp paired-end partial lane sequencing on a Novaseq X 25B flow cell with Novogene.

**Antibodies for CUT&Tag:**

| Target | Host Species | Target Species | Dilution | Manufacturer | Cat. No. | LOT# |
| --- | --- | --- | --- | --- | --- | --- |
| H3K4me1 | Rabbit | Mouse | 1:100 | Abcam | AB8895 | 1040931-12 |
| H3K27ac | Rabbit | Mouse | 1:100 | Abcam | AB4729 | 1086546-2 |
| H3K4me3 | Rabbit | Mouse | 1:100 | Millipore | 07-473 | 4103062 |
| H3K27me3 | Rabbit | Mouse | 1:100 | Cell Signalling Technologies | 9733 | 8, 27 |
| H3K9me3 | Rabbit | Mouse | 1:50 | Active Motif | 39162 | 23342141-12 |
| IgG | Guinea Pig | anti-rabbit | 1:100 | Antibodies-online | ABIN101961 | NE-200-042409 |

#### CUT&RUN

Cleavage Under Targets & Release Under Nuclease (CUT&RUN) was performed using the EpiCypher CUTANA CUT&RUN protocol following manufacturer instructions and using manufacturer recommended reagents, with minor modifications. Cells were differentiated on CellBIND dishes as previously described, and 500.000 cells were collected per antibody replicate, in duplicate per condition. Briefly, cells were washed with PBS, collected in accutase, neutralised with PBS, counted, and centrifuged for 5 minutes at 300 g and resuspended in Wash Buffer to a concentration of 1 million cells per ml, twice. Cells were bound to 1/10 volume (relative to cell suspension) BioMag Plus Concanavalin A beads (Cliniscience Ltd #86057-10) previously activated with Binding Buffer for 10 minutes at RT in DNA lo-bind Eppendorf tubes. Cells were subsequently incubated with 1:50 primary antibodies (including a negative IgG control) in Antibody buffer overnight at 4°C with gentle rocking. Beads were subsequently washed 3 times with 0.01% Digitonin buffer and resuspended in Digitonin buffer with pAG-MNase (EpiCypher CUTANA 15-1016). Tubes were placed on ice and 1 µL of 100 mM CaCl_2_ was added. Samples were incubated for 2 hours at 4°C with gentle rocking for targeted chromatin digestion. The reaction was stopped using Stop buffer which stops MNase enzymatic activity by chelating Ca^+^ ions. Tubes were then incubated for 10 minutes at 37°C to release cleaved chromatin and degrade RNA. DNA was purified using the NEB spin PCR DNA purification kit (NEB #T1130L). DNA concentration was quantified using a Qubit 2.0 Fluorometer. 5 ng of purified CUT&RUN-enriched DNA per sample was used to prepare Illumina NGS libraries following an adapted protocol with NEBNext Ultra II DNA library prep kit for Illumina (New England Biolabs #E7645S). Briefly, for NEBNext End prep, samples were incubated with NEBNext Ultra II Prep Enzyme Mix and Reaction Buffer. For adaptor ligation, unique adaptors were added to each sample, with NEBNext Ultra II Ligation Master Mix, Ligation Enhancer, and later USER enzyme. To clean up ligated DNA, 1.75 SPRIselect beads were used. DNA was then PCR amplified using NEBNext Ultra II Q5 Master Mix together with universal i5 and unique i7 primer from the NEBNext Multiplex Oligos for Illumina kit (New England Biolabs #E7335S). Post-PCR amplification involved a two-step SPRIselect bead cleanup at 0.8x followed by 1.2x to remove fragments larger than 350 bp and smaller than 150 bp. DNA was resuspended in 0.1x TE pH 8 buffer. Quality control was done using an Agilent D1000 ScreenTape (Agilent, # 5067-5582) on the Agilent TapeStation 4200 (JCBC NGS facility; Analysis Software Version 5.1) and a Qubit 2.0 Fluorometer. Pooled samples were submitted for 150 bp paired-end partial lane sequencing on a Novaseq X 25B flow cell with Novogene.

**Antibodies for CUT&RUN:**

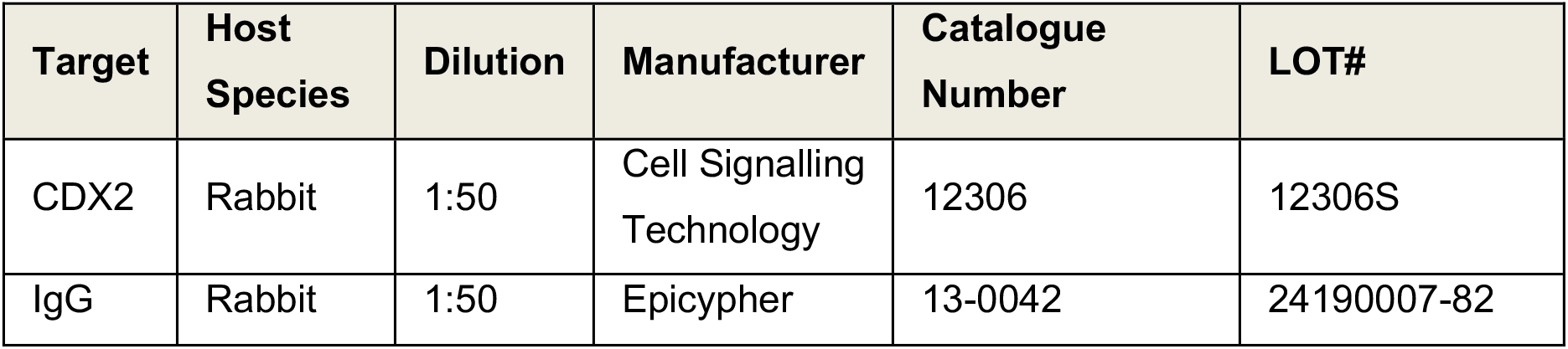

#### Bulk RNA-sequencing

Bulk RNA-sequencing was performed by collecting 1 million cells per sample in triplicates for RNA extraction. Briefly, cells were washed with PBS, harvested, centrifuged 5 mins at 300 g, washed in PBS and counted. The appropriate amount of cells was pelleted for 3 mins at 300 g and resuspended in 0.2x culture volume Trizol (Invitrogen, #15596018), and placed at −80 °C. RNA was extracted using the Direct-zol RNA Miniprep Plus Kit (Zymo Research, #R2072) following manufacturer instructions and stored at −80 °C. Quality control (QC) of total RNA was carried out using the Qubit™ RNA HS Assay Kit (Invitrogen, Q32852) and the RNA ScreenTape for Tapestation (Agilent, 5067-5576, 5067-5577, 5067-5578). Libraries were prepared using NEBNext® Poly(A) mRNA Magnetic Isolation Module (#E7490), NEBNext® Ultra™ II Directional RNA Library Prep Kit for Illumina® (#E7760), and NEBNext® Multiplex Oligos for Illumina® (96 Unique Dual Index Primer Pairs #E6440 #E6442) following manufacturer instructions. For QC of libraries, the Agilent D5000 ScreenTape System (Agilent, 5067-5588, 5067-5589, 5067-5590) was used. Libraries were pooled in equimolar ratios and submitted for 150 bp paired-end sequencing at the CRUK-CI Genomics Core Sequencing Facility and sequenced on a NovaSeq X 25B Flow Cell.

### 3. Data Analysis & Statistics

#### Quantitative immunofluorescence data analysis

CZI files were imported into FIJI (is just ImageJ, version 2.9.0/1.53t; Java 1.8.0_322 [64-bit]) and the same brightness and contrast settings were applied across all images for the same channels and the same laser settings. Average intensity z-projections with the same number of stacks across all cell types were saved as TIF files for further segmentation. Nuclear segmentation was carried out using CellProfiler (version 4.2.6) on the DAPI channel using the two-class Otsu thresholding method, and subsequent nuclear objects were used to quantify signal intensity across the other channels. Outputs included the original TIFs with an overlay of the nuclear objects, as well as csv files detailing the sample metadata and various quantifications of nuclear size, shape, signal intensity, etc. CSV files were subsequently imported into Rstudio (version 2024.04.1) for further statistical analyses and plotting.

Fluorescence intensity for each transcription factor was normalised to DAPI intensity at the single-cell level and log10-transformed. For each sample (≥50 cells/sample), the mean log10(DAPI-normalised intensity) was calculated. Sample means were aggregated by category (≥2 samples/category) within each experiment. To assess reproducibility of transcription factor expression trends across experiments, fold-changes were calculated relative to a fixed reference category within each experiment. For each category, the median of sample mean log10(DAPI-normalised intensity) values (≥2 samples/category) was used as the experiment-level value. Fold-change was computed for individual replicates against the mean of the reference condition within each experiment. Fold-change values from all experiments were pooled by category. Differences in fold-change distributions between categories were evaluated using Kruskal-Wallis tests, followed by Dunn’s post-hoc pairwise comparisons with Benjamini-Hochberg correction for multiple testing. Each plotted point represents one experiment-level fold-change.

| Sample | Size range (pixels) | Threshold strategy | Threshold method | Threshold smoothing scale | Bounds on threshold | Size of adaptive window |
| --- | --- | --- | --- | --- | --- | --- |
| EpiLC-CEpiLC | 40-110 | Adaptive | Otsu | Two-class | 0-1 | 140 |
| Mesoderm | 50-140 | Adaptive | Otsu | Two-class | 0-1 | 200 |
| SCP | 50-140 | Adaptive | Otsu | Two-class | 0-1 | 200 |

#### Single Molecule Localisation Microscopy data analysis

Data analysis of single-molecule localisations was carried out as previously described^46^ with minor changes. Briefly, single-molecule localisations were extracted from 3D DHPSF movies using easyDHPSF software^126^ in MATLAB (version 2023b) and the calibration file obtained prior to the imaging of each dataset. After manual inspection of different thresholds, a relative localisation threshold of 70 was set for all 6 angles of H2B datasets. Individual localisations were assembled into trajectories using ‘trajectory-analysis’ (https://github.com/TheLaueLab/trajectory-analysis). Molecules in subsequent frames within a 500 nm radius were considered part of the same trajectory and hence connected, with a frame gap tolerance of 2 frames, as previously described^44^. Additionally, the z-coordinate range of the true imaging signal was determined by calibration, and trajectories outside this range, corresponding to the beam light path, were excluded using the custom algorithm *dhpsf_localisation_cropper*. A custom algorithm developed in our lab called ‘analyze-tracking-data’ was then deployed, which uses a sliding window along each trajectory to output four biophysical parameters for each sub-trajectory – anomalous exponent (α), apparent diffusion coefficient (D_app_), length of confinement (L_c_) and drift magnitude^46^. Analysis was carried out using the following variable settings in the code: a sliding window of 9 frames for H2B-loci, 0.5 s exposure time, a minimum trajectory length of 4 frames. Since chromatin mobility can be classified into two subpopulations, slow- or fast-moving chromatin, we also set this algorithm to use a 4-parameter Gaussian mixture model, considering all four biophysical parameters (default), to classify sub-trajectories as either slow- or fast-moving and to output biophysical parameters separately for each trajectory type. The average parameter value for each replicate were plotted using custom scripts in RStudio (version 2024.4.1). To assess normality, we conducted the Shapiro-Wilk test and analyzed Q-Q plots before performing any comparisons. Based on the normality results, we used the Mann-Whitney-Wilcoxon test for non-parametric comparisons, applying the Bonferroni-Holm adjustment method for multiple testing corrections.

#### Quality control and adaptor trimming for FASTQ files

For all next-generation sequencing datasets (bulk RNA-seq, CUT&Tag, CUT&RUN, Micro-C), quality control and read trimming were performed on the raw FASTQ files. FastQC (v0.12.1) was used to assess sequence quality scores and adapter content, and results were aggregated using MultiQC (v1.27.1)^95^. Quality and adapter trimming were performed using Cutadapt (v5.0) using corresponding adapter sequences and parameters -q 20 -minimum-length 10^96^.

#### Micro-C data analysis

Micro-C data was pre-processed following the Dovetail genomics pipeline (https://micro-c.readthedocs.io/en/latest/). In brief, raw reads were trimmed using *cutadapt*^96^, and aligned using BWA-MEM2 (v2.2.1)^98^ and mm39 as reference genome with parameters −5SP -T0. *pairtools*^99^ and *pairix*^100^ were used to record valid ligation events and to index the pairs of genomic coordinates respectively to generate pair files. Reproducibility of the replicates was assessed at 100kb, 250kb, 500kb, 1Mb, 2Mb and 5Mb using HiCRep (v1.12.2)^101^. To detect significant interactions at higher resolution, pair files for each sample were combined and binned at 10 kb resolution using *cooler*^104^. Then, to reduce bias caused by differences in read depth, cool files were downsampled with respect to the dataset with the fewest intrachromosomal interactions. ICE-normalised binned matrices were generated using FAN-C(v0.9.28) on autosomes and the X chromosome^102^.

A/B compartment analysis was performed at 500kb resolution using *fanc compartment* with default parameters. For each chromosome, a Pearson correlation matrix was constructed, and the compartment type was assigned based on the first eigenvector. The sign of the eigenvector was corrected using average GC content derived from the mm39 reference genome. Pearson correlation matrices and corresponding eigenvector values were visualised using *fanc plot*. For A-B compartment shifting, each bin was assigned to A or B compartment state, and shifts were determined by comparing these bins to the preceding developmental stage. The percentage of bins in each shift category was plotted using *ggplot2* in Rstudio.

A/B compartment strength was assessed using cooltools and cooler^104,106^. The first cis eigenvector (E1) was calculated for each chromosome using *cooltools eigs_cis* with GC content calculated from GRCm39 (mm39) reference genome using *bioframe*. The orientation of E1 was determined by its correlation with GC content, and the eigenvector sign was inverted where necessary to ensure positive E1 values corresponding to GC-rich A compartments. Expected cis contact frequencies were calculated using *cooltools expected_cis*. Saddle plots were generated using *cooltools saddle* by partitioning genomic bins into 38 quantiles based on E1 values after excluding the lower and upper 2% of the eigenvector distribution. Observed-over-expected interaction frequencies between compartment bins were visualised as heatmaps using logarithmic colour scaling.

Topologically associated domain (TAD) boundaries were called at 50 kb resolution and window sizes of 600kb using *fanc insulation* and *fanc boundaries* modules. Insulation scores, TAD boundaries, and the interaction heatmaps were visualised using *fancplot*. TAD sizes were determined using the distances between the start and end of TAD boundaries. Number of TADs, TAD sizes, boundary strengths, and insulation scores at boundaries were visualised using ggplot2 in R. Statistical tests were performed using two-sided Wilcoxon ranked sum test and p-values were adjusted using the Benjamin-Hochberg procedure.

Statistically significant intra-chromosomal interactions were called at the binned contact matrices using FitHiC2 (v2.0.8)^61^, with extra settings -U 2000000 --L 40000 to constrain extreme interaction sizes. To focus on statistically significant and high-confidence interactions for downstream analysis, interactions called by FitHiC2 were filtered to retain those with false discovery rates (FDR) below 10^−5^ and minimum raw counts ≥ 10. Significant interactions were visualised using pyGenometracks (v3.9).

Subsets of chromatin loops with CREs or CDX2 binding at one or both anchors were identified using the *bedtools intersect* and *pairToBed* commands with the -u option^120^. CREs were defined by overlapping histone mark peaks using *bedtools intersect*, with the -f 0.25 option. Venn diagrams showing shared and unique CREs between conditions were generated using *intervene venn*.

Aggregate peak analysis (APA) was performed to visualise and compare contact signals for loops in each genomic feature or CDX2-binding category. Cool matrices were iteratively corrected using intrachromosomal contacts with *cooler balance*^104^. Then, the expected Micro-C signals for each chromosome were calculated using *cooltools expected-cis* command^106^. For each sample and category, pile-ups of observed over expected contact signals at the anchors of loop category were generated with *coolpup.py* using the balanced matrices^107^. Heatmaps of aggregated signals were plotted using *plotpup.py* with additional parameters *--flank 1000000* to minimize differences in background signals. The comparison of signals across samples or categories was performed using *dividepups.py* command.

#### Bulk RNA-seq data analysis

Quality-trimmed RNAseq reads were aligned to mouse mm39 primary genome assembly using STAR (v2.7.11b) with default settings^108,109^. BAM files were quality-checked using RSeQC (v5.0.4) and Qualimap (v2.3) to assess read distribution, gene body coverage profile, and GC content^110,111^. Read count was generated with Subread (v2.0.8) featureCount using Gencode M37 comprehensive gene annotation^109,117^. Principal component analysis was conducted and visualised using DESeq2 (v1.49.4)^113^.

Differential gene expression analysis was performed using edgeR (v4.7.3)^114^. Genes with low expression were subsequently removed using filterByExpr, and the count data was processed using the quasi-likelihood data analysis pipeline. Each cell type (CEpiLC, SM, SCP) was compared to the preceding differentiation stage to identify differentially expressed genes. Statistically significant differentially expressed genes were identified as genes with a false discovery rate (FDR) < 0.05 using *glmTreat* and an absolute log2 expression fold change (LFC) > 1. To visualise the data, raw gene counts of differentially expressed genes were converted to trimmed mean of M values (TMM)-normalised count per million (CPM) values using *edgeR*. The normalised gene expression was then converted to z-scores and visualised using *pheatmap* in R.

Gene ontology analysis was performed using *g:Profiler* R client (v0.2.3)^115^. For differentially expressed genes, all genes retained after edgeR *filterByExpr* were used as the custom background, and significant gene ontology terms were defined as entries with an adjusted p < 0.05. Redundant gene ontology terms were reduced using *rrvgo* R package (v1.20.0) with threshold = 0.7, and results were visualised using *rrvgo scatterPlot*^116^.

Gene set enrichment analysis was performed using Broad Institute Gene Set Enrichment Analysis (GSEA) desktop software (v4.4.0)^117^. Raw RNAseq gene counts were TMM-normalised using the CPM values from edgeR and adjusted to cls format for GSEA input. Gene set enrichment analysis was performed using mouse-ortholog hallmark gene sets (MH), curated gene sets (M2), and cell type signature gene sets (M8) obtained from the Molecular Signatures Database in GSEA software.

#### CUT&Tag data analysis

Reproducibility of CUT&Tag datasets were assessed using deepTools *plotCorrelation* with the parameter -c pearson and a bin size of 500 bp. To increase read depth, quality-trimmed CUT&Tag datasets were pooled, and aligned to mm39 primary genome assembly using BWA-MEM2 (v2.2.1) with default settings^98^. Following read alignment, SAM files were converted to BAM format, sorted by coordinates, and filtered using SAMtools (v1.21) *view* with parameters -F 2034 -f 2^118^. Duplicated reads were marked using Picard (v3.4.0). Reads overlapping suspected problematic regions were removed using BEDtools (v2.31.1)^120^. Peak calling was performed on the blacklisted BAM files using MACS3 (v3.0.3)^119^, which identified regions with histone mark enrichment. For H3K4me3 and H3K27ac marks, peaks were called using parameter settings --keep-dup all --nomodel --qvalue 0.05. For H3K4me1 and H3K27me3 marks, peaks were called with additional parameters -broad --broad-cutoff 0.1. Aligned reads and histone peaks were quality-checked using R package ChipQC (v1.44.0)^121^. The genomic distribution of MACS3 peaks across annotated genomic features was visualised using R package ChIPseeker (v1.44.0)^122^.

Differential peak analysis was performed to quantitatively evaluate CUT&Tag signals and detect genomic coordinates with cell-specific histone mark enrichment. Pairwise comparison of histone mark signals was performed using MAnorm (v1.3.0)^123^, using BAM files, read coordinates from MACS3 peak-calling, and additional parameters -p-cutoff 0.01 --m-cutoff 0.58 --window-size 2000 --summit-dis 500.

Active and primed enhancer regions were identified using CUT&Tag peaks from MACS3. Putative active enhancers were defined as the intersects of H3K27ac and H3K4me1 peaks. Primed enhancers were defined as regions with H3K4me1 but not H3K27ac peaks. Peak intersection was performed using BEDtools *intersect*^120^.

For visualisation, CUT&Tag signals were normalised and converted to BigWig format. Blacklist-removed BAM files were binned into 15 kb intervals, and the number of aligned reads per bin were obtained using Subread *featureCount* (v 2.0.8)^130^. Scaling factors were then calculated using CPM normalisation based on TMM-normalised effective library size in edgeR^131^. Normalised BigWig tracks were then generated from the BAM files and scaling factors using deepTools (v3.5.6) *bamCoverage*^125^, and visualised using Integrative Genomic Viewer (v2.19.2)^132^. Heatmaps of CUT&Tag signals at transcription start sites were visualised using deepTools *computeMatrix* and *plotHeatmap* using the normalised BigWig files and Gencode mouse M37 basic gene annotation^109^. Genomic coordinates from peak calling, differential analysis, and peak intersection were visualised using upset plot from Intervene (v0.6.5)^97^.

#### CUT&RUN data analysis

CUT&RUN datasets were analysed using similar methods to the CUT&Tag. Quality-trimmed reads from each replicate were aligned using BWA-MEM2 (v2.2.1)^98^, and unpaired reads, supplementary alignments, and reads overlapping CUT&RUN suspected problematic regions were filtered out. Peak calling was performed using MACS3 (v3.0.3) and additional parameters --keep-dup all --nomodel --qvalue 0.05^119^. Quality checking and reproducibility analysis on aligned reads and peaks were performed using ChipQC (v1.44.0)^121^. For each sample, consensus binding sites were defined as the intersections from both replicates using BEDtools *intersect*^120^.

CUT&RUN signals were normalised using background normalisation as described for the CUT&Tag analysis. Each replicate was converted into a normalised BigWig track. For each cell type, the mean signal across replicates was calculated using Ensembl WiggleTools (v1.2.1)^124^ and converted to BigWig format for visualisation.

CDX2 CUT&RUN peaks were assigned to genes using the peak-to-TSS annotation. Duplicate gene–peak pairs arising from multiple transcript or TSS annotations were removed before gene-level quantification, and CDX2 binding strength was calculated as the sum of the scores of all unique CDX2 peaks assigned to each gene.

RNA expression was quantified as CPM using effective library sizes adjusted by TMM normalisation factors. Genes were retained if CPM was at least 1 in at least two of three biological replicates in either condition. Expression values were transformed as log_2_(CPM + 0.5) for each replicate, and expression change was calculated as the difference between mean transformed expression in CEpiLC and EpiLC.

Expression-change values were matched to CDX2 binding scores by Ensembl gene identifier. CDX2 binding scores were log_10_-transformed for visualisation, and the association between CDX2 binding strength and transcriptional change was assessed using a two-sided Spearman rank correlation. For annotation in the scatter plot, genes were labelled when they fell within both the top 10% of summed CDX2 binding scores and the top 10% of absolute expression changes. Analyses and visualisation were performed in R using *tidyverse* and *ggrepel*.

#### Schematic and graphical representations

Statistical analysis and figure generation was performed in RStudio unless stated otherwise. Schematic diagrams were generated using BioRender.com(Wurmser, A. (2027) https://BioRender.com/e1hlesa and https://BioRender.com/y2i2358). Panels were assembled using Adobe Illustrator.

### 4. Chromatin modelling using a generalised random-crosslinked (RCL) polymer model

We represented chromatin organisation based on our previous randomly cross-linked (RCL) polymer model^133^. The RCL polymer consists of *N* beads, with positions *R* = (*r*_1_, …, *r_N_*), consecutively connected by harmonic springs to form a Rouse-like backbone^27,134^. Chromatin loops were formed by additional fixed harmonic springs between non-consecutive beads. To model a TAD, we also connected the first and last beads to form a closed polymer chain. No bending potential or volume exclusion were included. The total potential energy of the polymer is given by

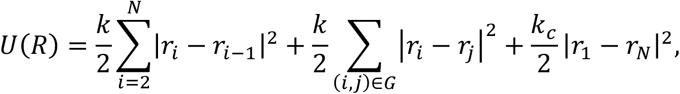

where *G* is the set of loop-forming bead pairs; *k* and *k_c_* are the spring constants. The dynamics of each bead *i* in the polymer were described by the Langevin’s equation:

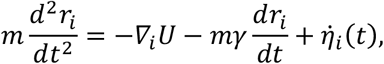

where *m* is the bead mass, *γ* is the friction coefficient, and *η̇_i_*(*t*) is the random thermal force acting on each bead (Brownian motion).

Numerical simulations were performed at constant temperature using molecular dynamics with implicit solvent in LAMMPS (29 Aug 2024 version)^135^. Simulations used Lennard–Jones (LJ) reduced units, with bead mass *m* = 1, characteristic length *σ* = 1, and energy scale *ε* = *k_B_T* = 1. The corresponding characteristic time is 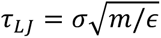. The Langevin equation was integrated using the velocity-Verlet algorithm with a local friction of 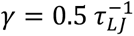 and an integration timestep of *dt* = 0.012 *τ_LJ_*. Simulations were performed in a cubic box of side length 400 *σ* with non-periodic boundaries. Each polymer was equilibrated for 1 × 10^6^ integration steps, followed by a production run of 2 × 10^5^ steps. Polymer coordinates were recorded every 1,000 production steps, corresponding to a sampling interval of *Δt* = 12 *τ_LJ_*.

We set *N* = 500 beads to account for the size of a TAD, with each bead representing 1kb of chromatin. We set *k* = 1.0 *ε*/*σ*^2^ for backbone and loop-forming springs, and *k_c_* = 0.5 *ε*/*σ*^2^ for the weaker spring closing the chain. To understand the effects of short and long loops on chromatin mobility, we varied the sets of loop-forming springs and simulated three loop simulation:

1. The **“Mixed”** simulation contained three evenly spaced long loops of 150 kb and 12 evenly spaced short loops of 30 kb overlapping;
2. The **“Δlong”** simulation removed the long loops and contained only the 12 short 30-kb loops;
3. The **“Δshort”** simulation contained only the three long 150-kb loops. This gives the following sets of loop-forming bead pairs for each simulation:

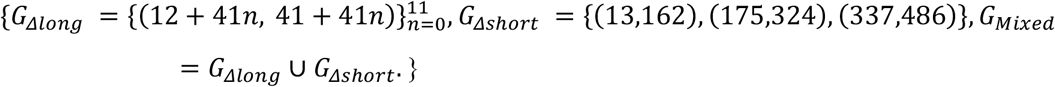

For each simulation, 50 independent realisations were simulated.

To quantify chromatin mobility, we computed the apparent diffusion coefficient (D_app_), confinement length (Lc) and anomalous exponent (*α*), following the single particle trajectory analysis framework described above^46^.

To compute the encounter probability, we define a contact when the spatial distance between two beads is less than 1.5 *σ*, i.e. 1.5 bead (characteristic length). Contact frequency between bead *i* and bead *i* + *s* (*C_i_*_,*i*+*s*_) was calculated as the fraction of frames satisfying this criterion, pooling all recorded frames across 50 realisations per simulation. Encounter probability at bead separation *s* was given by taking the mean of contact frequencies:

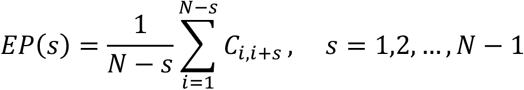

where *N* is the number of beads.

## Author contributions

**Conceptualisation:** AAMW and SB with help of BS. **Funding acquisition:** AAMW and SB. **Project administration:** SB. **Supervision:** SS, TEB and SB. **Investigation:** (SMLM) AAMW with help from GGA and MJ. (Micro-C, RNA-seq, CUT&RUN) AAMW. (CUT&Tag) AAMW with help from MMP. (Quantitative immunofluorescence) AAMW. (Polymer simulations) YC and DH. **Formal Analysis:** (Micro-C, RNA-seq, CUT&Tag and CUT&RUN) LM, NM with help from AAMW and NdeW. (qIF, SMLM) AAMW. **Validation:** AAMW. **Data curation**: AAMW and SB. **Visualization:** (Micro-C, RNA-seq, CUT&Tag, CUT&RUN) AAMW, LM and NM. (SMLM, qIF) AAMW. **Resources:** (Cell lines expressing HaloTag-H2B) AAMW with help from MJ. (SMLM instrument building and maintenance) ZZ and DK. **Methodology:** No new methods. **Software:** No new software. **Writing – original draft:** AAMW and SB. **Writing – review & editing:** TEB with comments from all authors.

## Acknowledgements

We thank all the members of the Basu lab for insightful and critical discussions, especially Liviu Pirvan for his early analysis of the Micro-C datasets, and Ben Steventon for critical reading of the manuscript. We thank the Cambridge Stem Cell Institute (CSCI) Genomics Core Facility and Cancer Research UK Cambridge Research Institute for quality control of RNA-seq libraries and Illumina sequencing. We thank Darran Clements in the CSCI Imaging Facility for training students and assisting with experiments. We thank the Cambridge BRC Phenotyping Hub for support with flow cytometry. We thank Sally Lees and the CSCI Tissue Culture Core Facility for technical advice and support. We thank Brian Hendrich for his kind gift of the PBase plasmid. We thank Despina Stamataki and James Briscoe for the kind gift of the *Cdx1/2/4* knockout embryonic stem cell line.

AAMW was funded by the Wellcome Trust PhD programme in Stem Cell Biology and Medicine (218481/Z/19/Z). GGA was funded by the UKRI Biotechnology and Biological Sciences Research Council (BB/W000423/1, BB/W000423/2). SB was funded by the Wellcome Trust (203151/Z/16/Z), UKRI Medical Research Council (MC_PC_17230), Trinity College Stem-Cell Medicine Senior Postdoctoral Researcher Fund and Imperial College London. ZZ was funded by the UKRI Engineering and Physical Sciences Research Council (EPSRC) (EP/W015005/1). TEB is generously supported by the Wellcome Trust (WT G117980) and the Cambridge Stem Cell Institute. MMP was funded by the UKRI Medical Research Council (MR/X008479/1).

## Notes

### Competing Interest Statement

The authors have declared no competing interest.

